# Exploring subacute effects of a DMT and harmine meditation retreat on the brain’s control energy landscape

**DOI:** 10.64898/2026.11.07.764538

**Authors:** Isabella Llano Aristizabal, Kat Hefter, Suman S. Kulkarni, Klemens Egger, Milan Scheidegger, Diana I. Tamir, Dani S. Bassett

## Abstract

Modeling the brain as a controllable structural network allows the estimation of the control energy (CE) required to reach functional states, where low-CE states are more accessible. Psychedelics are thought to acutely flatten the brain’s CE landscape, broadening the repertoire of reachable states, but whether this reorganization persists into the subacute “afterglow” window is largely unknown. This exploratory study uses rs-fMRI from a randomized, double-blind, placebo-controlled trial (N = 39; 19 verum [N,N-dimethyltryptamine (DMT) + harmine, Ayahuasca-inspired formulation], 20 placebo) scanned before and after a three-day meditation retreat. We compute CE on a group human connectome project (HCP) structural connectome under two approaches that emphasize long and short timescale dynamics. Globally, we found no systematic shift in the distribution of CE between transitions, however mean CE increased in both groups at the long timescale approach. Surviving network effects emerge when a network’s intrinsic timescale matches the stabilization regime. In the long timescale regime, the central executive network (CEN) showed an increase in post-retreat CE within the verum group that survived Benjamini-Hochberg False Discovery Rate (FDR-BH) correction. Although CE and functional connectivity are distinct, this higher CEN cost may relate to the reduced within-network connectivity previously reported subacutely after psilocybin. The default mode network (DMN) also showed a post-retreat CE increase in both groups, but neither change survived FDR correction. In the short timescale approach, the visual network (VN) showed a FDR-BH CE decrease within the verum group, consistent with reduced dynamical demand in a region acutely engaged by DMT. To examine whether changes were concentrated in less accessible states, we additionally analyzed rare brain states, defined by the top decile of information content. This analysis revealed a modest session × condition interaction in the long timescale approach. Overall, the two timescale lenses surface different network effects, that align with cortical timescale hierarchies. These findings point to network- and timescale-specific, rather than global, CE changes in the afterglow window, though effects are small and heterogeneous and require confirmation in larger samples. Characterizing whether and where psychedelic effects persist into the subacute window is relevant to psychedelic-assisted therapy, including safety, therapeutic timing, and durability of drug effects. More broadly, considering cortical timescale hierarchies may help identify network-specific CE changes that might be less apparent when networks are evaluated under a single dynamical timescale.

**Significance Statement:** Serotonergic psychedelics are theorized to increase flexibility facilitating transitions between functional states. Control energy quantifies the energetic cost of these transitions given brain structure, defining a control energy landscape over functional states. Recent studies show a flattening of this land-scape under psychedelics. Whether it remains altered after acute effects is unclear. Using fMRI from a randomized, double-blind meditation retreat comparing placebo with an Ayahuasca-inspired formulation, we find that control energy largely returns to baseline in both groups, with subtle, timescale- and network-specific effects that do not produce large, lasting alterations detectable at rest. At long timescales, control energy increased in the central executive network, while it decreased in the visual network at shorter timescales, in the psychedelic group. This is important for the safety, interpretation, and therapeutic use of psychedelics.

---

**I**n recent years, serotonergic psychedelics or entheogens (e.g., psilocybin, mescaline, LSD, and N,N-dimethyltryptamine (DMT)), along with other inducers of non-ordinary states of consciousness like MDMA and meditation, have been investigated for their therapeutic potential across a wide array of psychiatric conditions (1– 5). These interventions are characterized by their ability to generate transient alterations of consciousness driven by changes in perception, mood, cognition, and self-awareness, often leading to “mystical” or “peak” experiences (6, 7).

Neurobiologically, serotonergic psychedelics have been found to act as agonists of the serotonin 5-*HT*_2*A*_ receptor (8). This effect has been proposed as a mechanism of “disintegration” of the intrinsic functional network structure, potentially broadening the repertoire of available brain states (9). Network-based analyses have frequently reported reduced within- and increased between-network connectivity (10–13). On this point, a recent mega-analysis found increased connectivity between transmodal associative and unimodal sensorimotor networks as the strongest and most consistent effect. On the other hand, within network connectivity reductions were more variable overall and most consistent in sensorimotor networks over transmodal ones (14). Decreased modularity has also been reported after psilocybin treatment and has been associated with improvements in depressive symptomatology (9). Further, changes in functional connectivity have been reported to correlate with the intensity of the subjective experience, highlighting inter-individual variability in the effects of psychedelics (13, 15). Psychedelic researchers have proposed the RElaxed Beliefs Under Psychedelics (REBUS) model (16). It postulates that psychedelics, produce a decreased precision-weighting on prior beliefs, which then loosens high-level constraints on perceptual and cognitive processing (16). Increased brain entropy and reduced energetic barriers between states emerge as features of this framework (16, 17).

Network control theory (NCT) provides a quantitative method to assess the ease with which the brain can move through functional states given a specific structural network (18–21). Within this framework, control energy (CE) quantifies the minimum input energy required to drive the system from one functional state to another, given its underlying structure. Calculating CE across a set of transitions creates a distribution of transition costs that characterizes the brain’s control energy landscape. Previous work using NCT to study the dynamics of the resting-state brain has shown a reduced CE landscape under serotonergic psychedelics including LSD, psilocybin (22) and DMT (23), consistent with theoretical models like REBUS that suggest that psychedelics make transitions between brain states more accessible. Prior work has also found that state transitions with high control energy requirements are less frequent (24, 25). Since psychedelics seem to reduce the CE needed to make state transitions and flatten the energy landscape (22, 23), we hypothesize that psychedelics might affect rare high-energy states differently. This would be consistent with reports of an expanded repertoire of states during the psychedelic state (26).

Most studies supporting the REBUS framework come from imaging during the acute state (27). Subacute effects, defined as those occurring from 1 day to 1 month after using a psychedelic (28), have been consistently reported anecdotally but have only recently become the subject of formal study. A meta-analysis of subacute effects found reports of reduced psychopathological symptoms, increased well-being, and beneficial changes in perception of self, others, and the environment, along with some reports of mild to severe adverse effects (e.g., headaches, sleep disturbances, and psychological distress in some individuals) (28). It has been hypothesized that these “subacute effects” might enhance responsiveness to psychotherapeutic interventions. Despite this potential, the “psychedelic afterglow” has only one recent validated measurement tool (28, 29). Neuroimaging studies of this period remain limited with a specific focus on connectivity. Two subacute studies of psilocybin have reported: decreased functional connectivity in the central executive network (CEN) 1 week post-treatment in healthy volunteers (30), and increased functional connectivity within the default mode network (DMN) 1 day post-treatment in patients with treatment-resistant depression (31). A study of Ayahuasca reported increased connectivity within the salience network, decreased connectivity within the DMN, and increased connectivity between the salience network and DMN one day after administration (32). However, control energy effects in this subacute window have not been reported.

Additionally, recent efforts have begun to explore the interactions and synergies between psychedelic therapy and meditation (7, 36). To this end, Egger & Meling *et al*. conducted a randomized resting-state fMRI study in which 40 participants were scanned the day before and after a three day meditation retreat, where half consumed an Ayahuasca inspired formulation (DMT + harmine), and the other half a placebo (6, 37). Their analyses revealed mixed functional connectivity outcomes: meditation alone decreased functional connectivity between parts of the salience (SN), default mode (DMN), visual (VN), attention, and frontoparietal (CEN) networks. In contrast, DMT + harmine mostly increased between-network connectivity between VN–SN and, more weakly, between VN–DAN. The authors found no sustained alterations in cortical gradients post-intervention, suggesting that large scale hierarchical organization returns to baseline shortly after the experience (6, 37). This finding raises the possibility that subacute effects may emerge at more local or network-specific levels rather than through global reorganization.

Importantly, brain networks have been shown to operate on different timescales of temporal dynamics. Research has shown unimodal sensory regions tend to have faster dynamics, whereas transmodal associative regions, tend to operate over slower timescales (38–40). Given this temporal hierarchy, we also considered whether network control energy effects could vary according to the temporal properties of the networks involved.

Building on the findings of Meling & Egger et al., the present study performs an exploratory analysis to evaluate possible changes in control energy in participants who underwent meditation alone (placebo) or meditation augmented by DMT + harmine (verum) in a three day retreat. To account for potential timescale differences we compute CE under two stabilization schemes, one that enforces decay across all modes, emphasizing shorter timescales and another that allows for more persistent activity, emphasizing longer timescales. These regimes represent different modeling assumptions with complementary perspectives on network dynamics, rather than direct measures of intrinsic cortical timescales. Given the pre- and post-intervention nature of the data, we expect subtle effects, with possibly larger differences in the default, salience, and control networks (DMN, SN, CEN) and in the verum group, consistent with prior work on the triple network model in psychopathology (41) and the therapeutic effect of psychedelics (9). This analysis provides insight into the possibility of control energy changes after psychedelic augmented meditation, thereby increasing our understanding of subacute effects of psychedelics, which can help orient their therapeutic capabilities towards impactful and grounded interventions.

## Results

Our analysis was structured to detect potential subacute effects of psychedelics or psychedelic augmented meditation across different scales of resolution. We first asked looked at CE changes per transition, where each participant has a distribution of values, and also more broadly, changes in mean CE between groups. We then examined whether any changes localize to specific functional networks, motivated by the observation that meditation and psychedelic effects have been shown to affect transmodal associative and unimodal sensory networks differently (9, 14, 31, 42). Finally, because previous hypotheses and research have talked about a widened repertoir of functional states after psychedelics (26) and a flattened CE energy landscape (22, 23), we asked whether the energy landscape governing rare (high information content) brain states was specially affected.

As stated, we presented results emphasizing long and short timescale dynamics. The formal reasoning of the timescale interpretation of these stabilization schemes is in Methods and in Supplementary Information (SI) Text for interested readers. To avoid confusion with intrinsic cortical timescales we refer to our stabilization modeling choices as long and short timescales, whereas intrinsic activity will be referred to as slow and fast. Main text results use the Schaefer 200 parcellation; the 100 and 400 parcellations are reported in the SI for robustness. Given the nature of the data and the absence of sustained large scale reorganization previously reported in this dataset (6), we expect surviving effects to be modest, localized, and likely heterogeneous.

### Overall distribution of control costs for typical transitions is largely stable, with timescale-specific changes in mean control energy

To assess whether control energy (CE) shows an overall sustained change following the retreat, we first asked wether each participants distribution of transition-level CEs had changed between sessions. Findings of a flattened CE landscape during psychedelics motivated this question. We reasoned that a subacute effect of this reorganization could manifest as a redistribution or broadening of the distribution of energy costs for typical transitions rather than just in mean CE. For this we calculated mean CE across all time transitions for each participant in both the short and long timescale regimes and compared each participant’s pre- and post-retreat distributions of whole-brain transition energies, using two-sample Kolmogorov–Smirnov. We found no systematic shifts in CE distributions, significant differences occurred only in a small subset of individuals (Long timescale: 10/39, 7 after FDR-BH correction. Short timescale: 12/39, 4 after FDR-BH correction) and this did not relate to what group they belonged to (SI Fig. S1).

Because this comparison only tested whether each participants CE distribution from session 1 and session 2 came from the same probability distribution, we also analyzed changes in mean CE. To this end, we calculated each participant’s change in CE between sessions (Δ*CE* = Session 2*−* Session 1) and tested each group’s change against zero (Wilcoxon signed-rank *W*, FDR-BH across the two groups) and the difference between groups (Mann–Whitney *U*). Individual trajectories showed substantial variability in global mean CE under both timescale regimes (Figure 2 *A-B*), while the corresponding change scores illustrate the distribution of these individual changes (Figure 2 *C-D*). In the long timescale regime, CE increased on average in both groups, with significant within group increases relative to baseline for both placebo (*W* = 51, *p*_FDR_ = 0.044, *d*_*z*_ = 0.46) and verum (*W* = 38, *p*_FDR_ = 0.041, *d*_*z*_ = 0.60). However, groups did not differ significantly from each other (*U* = 207 *p* = 0.643, *d* = 0.03) (Figure 2*C*). These results were preserved for the 400, but not 100, parcellation (SI Appendix Fig. S2). In the short timescale regime, both groups showed a non-significant decrease relative to baseline (placebo: *W* = 66, *p*_FDR_ = 0.307, *d*_*z*_ = *−*0.28; verum: *W* = 85, *p*_FDR_ = 0.709, *d*_*z*_ = *−*0.14), with no difference between groups (*U* = 152, *p* = 0.292, *d* = *−*0.19)(Figure 2*D*). This direction of findings is preserved across all three parcellations (SI Appendix Fig. S3). Together, these results indicate that there was no systematic redistribution of energy costs to transition between states. However, mean CE did change across sessions with effects being significant and shared at the long timescale regime and non-significant at the short timescale regime. The substantial variability and the regime-dependent direction of changes in mean CE motivated our approach at the network level where we consider CE contributions to a transition per node.

**Fig. 1.**
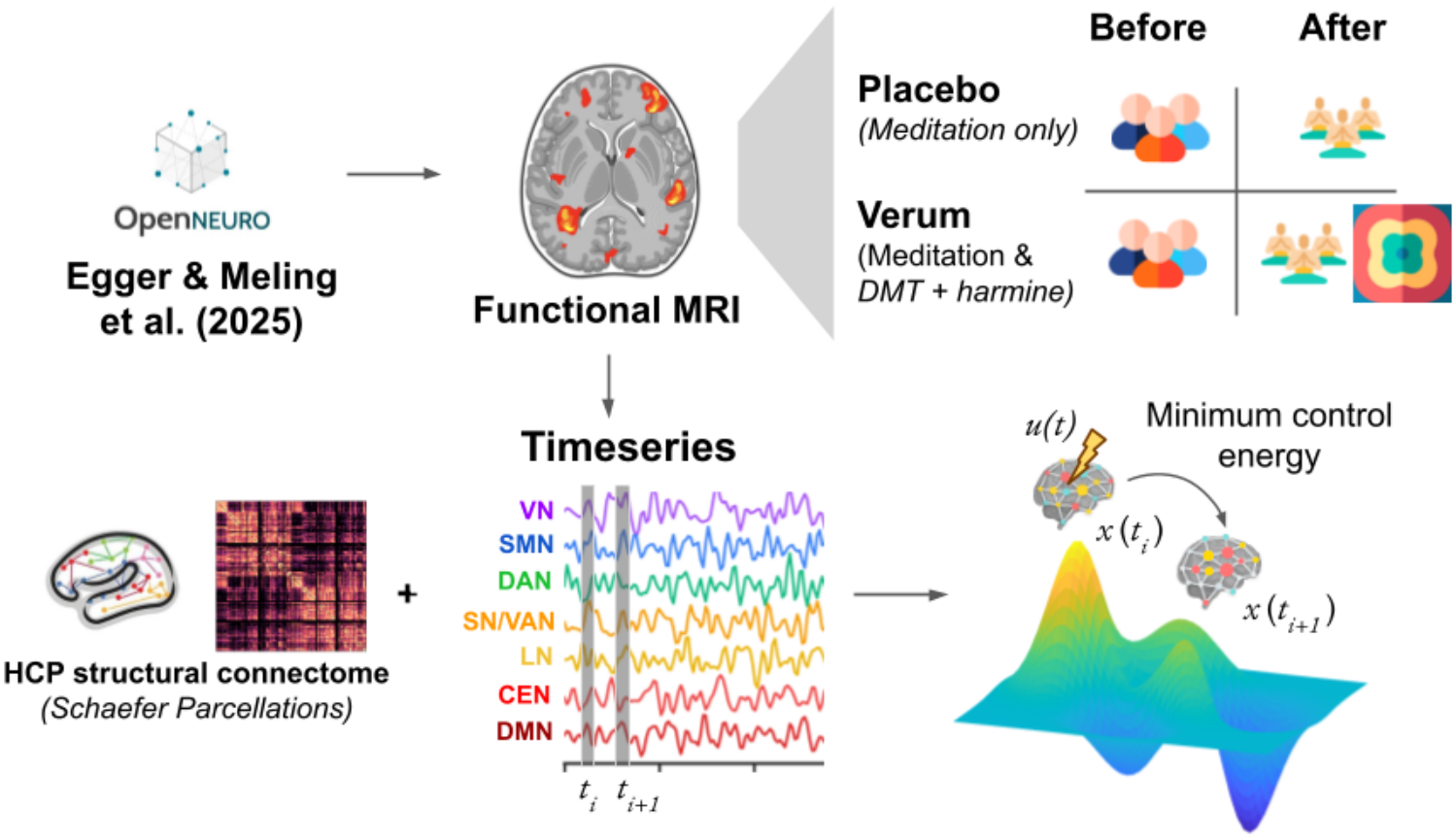
Study design and network control theory pipeline. rs-fMRI data was obtained from a randomized, double-blind, placebo-controlled study (33) in which participants were scanned before and after a three-day meditation retreat under either placebo (meditation only, *n* = 20) or verum (meditation + DMT + harmine, *n* = 19) conditions. BOLD time series were extracted following the Schaefer (34) brain parcellations (100, 200 and 400 regions) where each region belongs to one of seven networks: visual (VN), somatomotor (SMN), dorsal attention (DAN), salience/ventral attention (SN/VAN), limbic (LN), central executive (CEN), and default mode (DMN) (35). Using a group-averaged structural connectome from the Human Connectome Project (HCP) as the system matrix *A*, we compute the minimum control energy *u*(*t*) required to drive the brain from each consecutive state *x*(*t*_*i*_) to *x*(*t*_*i*+1_) across the entire time series, yielding a control energy contribution for every transition.

**Fig. 2.**
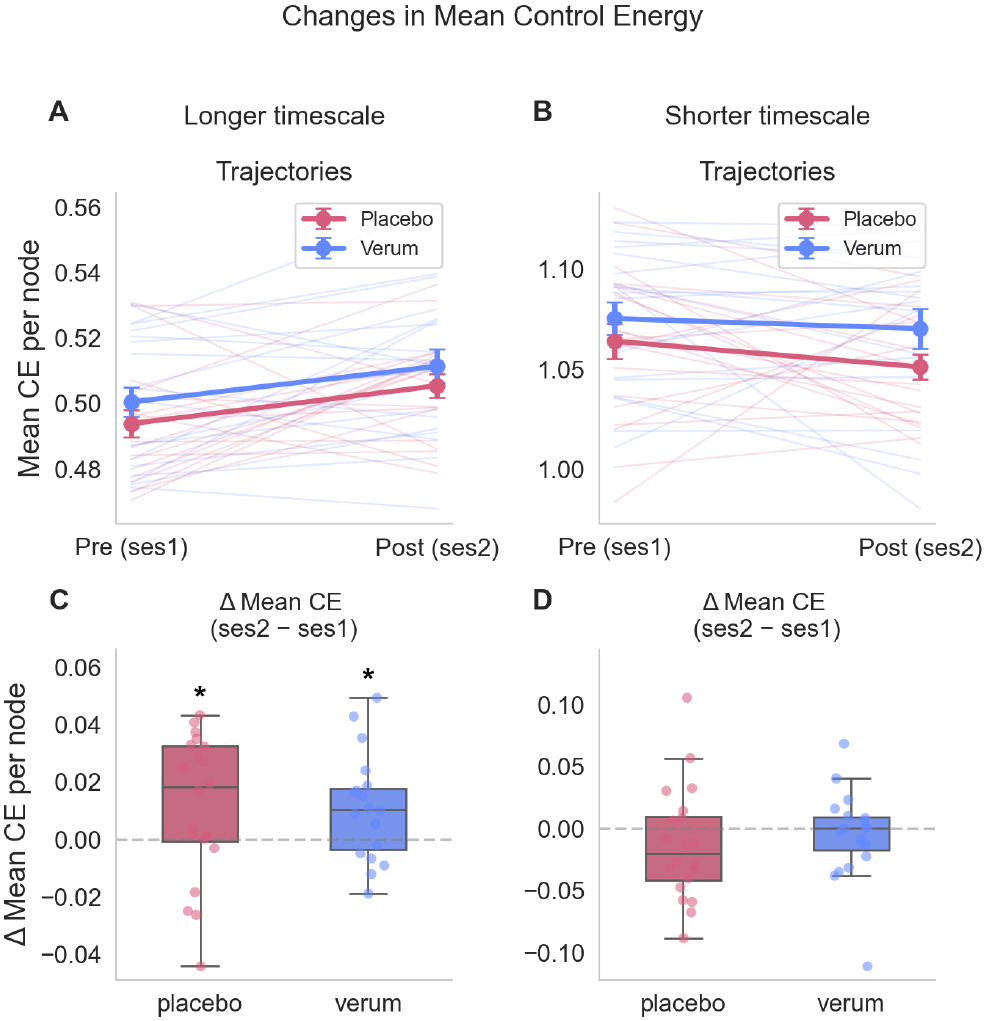
Changes in mean control energy across sessions and condition. *(A-B)* Individual subject trajectories (faded lines) and group means for placebo and verum participants from Session 1 (pre-retreat) to Session 2 (post-retreat), in the long and short timescale regimes, respectively. *(C-D)* The corresponding distributions of individual change scores (ΔCE = Session 2 *−*Session 1). Significance markers above boxplots represent within-group Wilcoxon signed-rank test (Δ CE vs. 0) where * = FDR-BH corrected *p <* 0.05. See SI Fig. S3 and S4 for results on 100 and 400 parcellations.

### Network effects are timescale specific and align with intrinsic cortical dynamics

To provide a general overview, the previous analysis aggregated node control energy and explored the mean changes between groups and distribution changes within participants. Because psychedelics have been shown to have network specific effects (14) and because networks have different intrinsic dynamics (39), we also sought to explore effects at the network level. For each participant and session, we averaged control energy across all state transitions to obtain one CE value for each node, and then averaged nodes within each Yeo-7 network. We then calculated the change from Session 1 to Session 2 for each network and tested these changes within each condition using Wilcoxon signed-rank tests, with FDR-BH correction across the 14 network *×*condition tests. At long timescales, CE increased in the central executive network (CEN) in the verum group (*W* = 23 *p*_FDR_ = 0.034, *d*_*z*_ = 0.83) with the same effect observed at the finer parcellation (SI Fig. S4). The Default Mode Network (DMN) also showed nominal CE increases in both groups (verum: *d*_*z*_ = 0.69, *W* = 33, *p* = 0.011 and placebo: *d*_*z*_ = 0.60, *W* = 43, *p* = 0.019) (Fig. 3*A*), the direction and magnitude of this trend was present at all parcellations but did not survive correction in any of them (SI Fig. S4). In the short timescale regime, only the Visual Network (VN) in the verum group showed a significant post-retreat decrease in CE (*W* = 22, *p*_FDR_ = 0.028, *d*_*z*_ = *−*0.91) (Fig. 3B). This result was the largest and was consistent across all three parcellations (SI Fig. S5). No between-group differences reached significance in either timescale regime. These findings showed CE changes localized to specific network and timescale combinations which then motivated subsequent exploration of whether effects might also be concentrated in particular functional states.

**Fig. 3.**
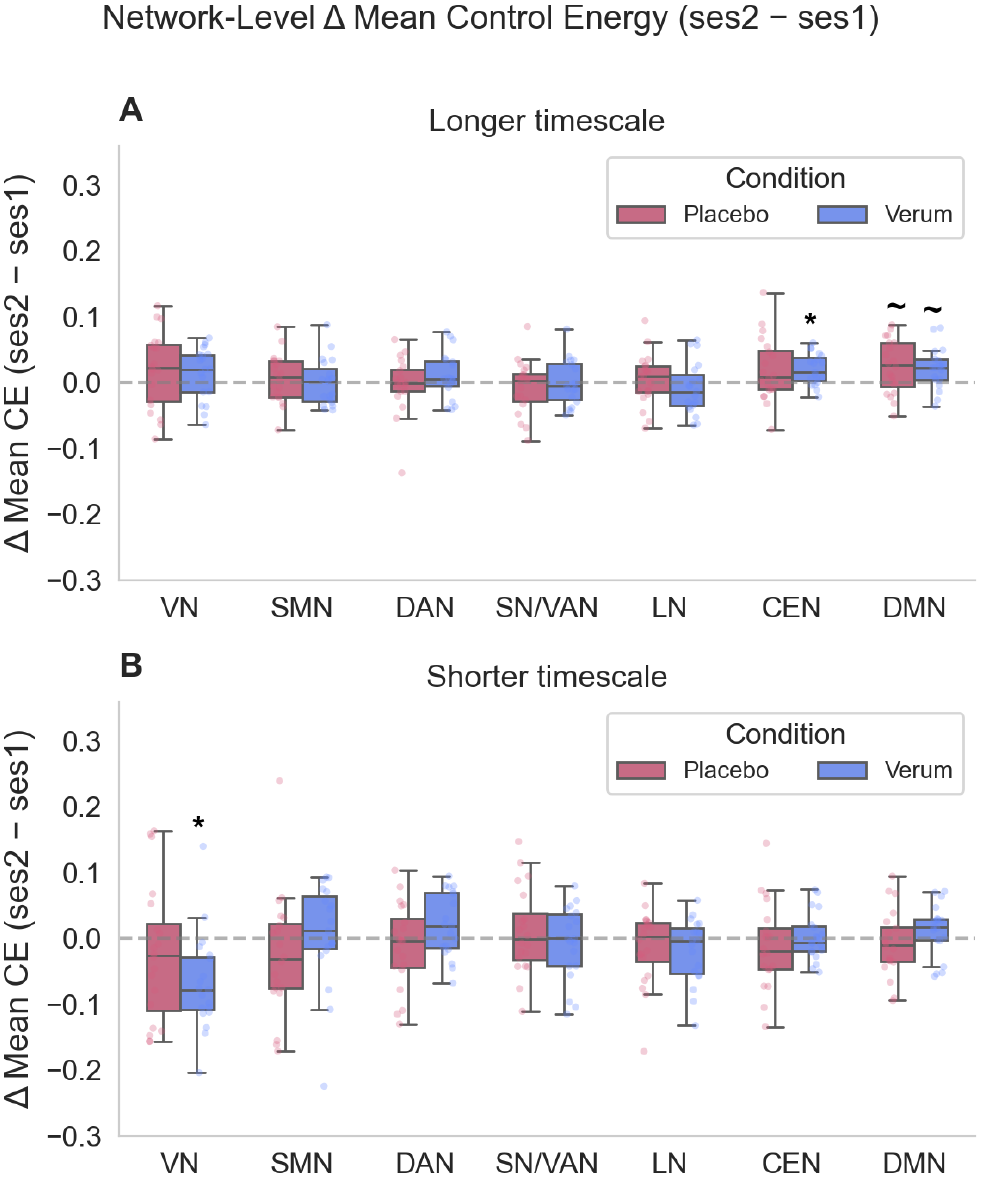
Changes in mean control energy at the network level. Δ Mean CE (Session 2 - Session 1) is shown for each of the canonical Yeo-7 functional networks for placebo (pink) and verum (blue) participants. Networks are ordered according to their approximate intrinsic timescales where motor and visual networks are fast responding and associative, higher order networks are slow responding. Boxplots display the distributions across participants, individual data points are overlaid. Markers above condition boxes reflect within-group Wilcoxon signed-rank tests (Δ CE vs. 0): * = *p*_FDR_ *<* 0.05 while *∼* = *p <* 0.05 . *(A) Results in the long timescale regime*. Here, the Central Executive Network (CEN) showed a significant post-retreat change in the verum group (*p*_FDR_ = 0.034), and the Default Mode Network (DMN) showed uncorrected trends in both conditions (see SI Fig. S4). *(B) Results in the short timescale regime*. Here, the Visual Network (VN) has a significant post-retreat decrease in the verum group only (*p*_FDR_ = 0.028.)

### Rare state specific energy landscape is affected differently across timescales

Taken together, the absence of systematic distribution changes and the presence of localized, timescale-specific network effects led us to ask whether residual subacute effects might also be concentrated in specific functional states. This is particularly relevant because the flattening of the energy landscape due to psychedelics has been proposed to facilitate access to otherwise hard-to-reach states. To explore this possibility, we used information content (IC) as a measure of state rarity, where rare states have higher IC and more frequent states have lower IC (25). Because IC is defined per state, and CE is defined per transition, we related each state’s IC to the CE required to enter (CE_in_) and leave it (CE_out_). IC is continuous, but representing all states simultaneously is visually uninformative, so we grouped states into three IC tiers representing high, middle, and low rarity as discrete samples along the continuum. When considering brain regions averaged over subjects, rare states, had the highest total energy regardless of timescale, condition or session followed by middle and frequent states (Figure 4). However, the relative cost of entering versus leaving a state depended on the stabilization regime used to interrogate the dynamics. In the long timescale regime, rare states required more energy to exit than to enter (Figure 4 A-D) whereas, in the short timescale regime, rare states required more energy to enter than to exit (Figure 4 E-H). This reversal follows from the different dynamical constraints imposed by the two stabilization regimes. The short timescale regime constrains the model to decaying activity in the absence of input, such that reaching rare, high-energy states requires constant input against this dissipation. In contrast, the long timescale regime permits the coexistence of decaying and expansive eigenmodes, allowing activity to persist. A reduced pull towards dissipation resulted in a higher relative cost of leaving an established high-energy state. This stabilization effect is also evident in our observation of near identical relationships before and after the retreat and in placebo and verum groups. Differences are dominated by timescale regime. Thus, the two regimes provide complementary views of how the entry and exit costs of states depend on the assumed dynamics of the model.

**Fig. 4.**
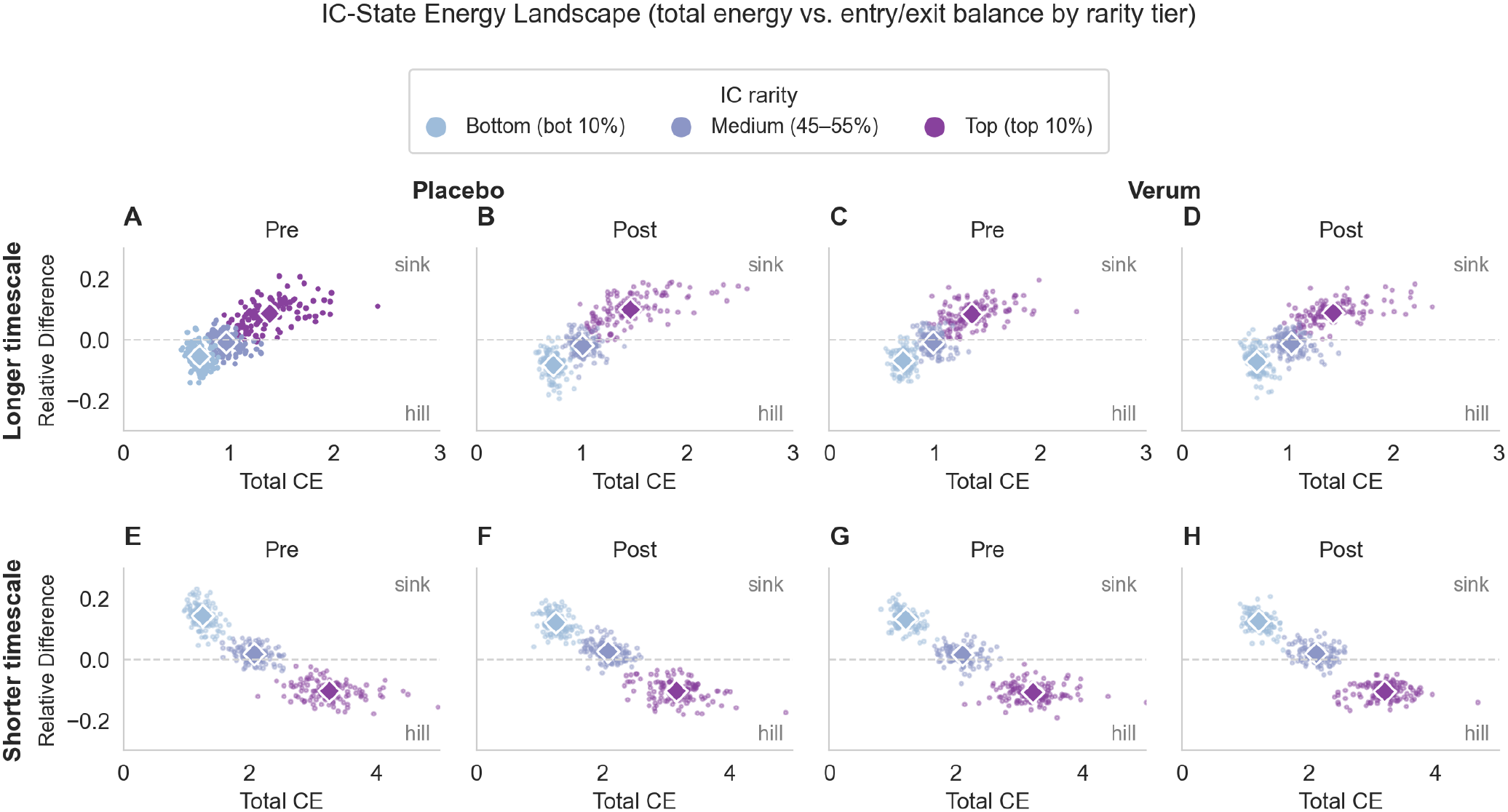
Control energy landscape across states of different rarity. We calculated information content (IC) for each state as a measure of state rarity, where higher IC indicates a rarer state. IC is a continuous quantity but we showed here three discrete samples along the continuum. States are grouped into high (top decile, purple), medium (45th–55th percentile, sky blue), and low (bottom decile, light blue) IC groups. For each state, we plotted total control energy CE_total_ = CE_in_ + CE_out_, which measures how costly a state is to visit, against the relative difference between the energy required to enter and leave the state Δ_rel_ = (CE_out_*−* CE_in_)*/*CE_total_. The dashed horizontal line marks equal entry and exit energy cost Δ_rel_ = 0. States above the line are more costly to leave than to enter (“sinks”), while states below are more costly to enter than to leave (“hills”). Each point represents a brain region averaged across subjects and diamonds indicate the mean of each IC group. *(A–D)* Results for the long timescale regime. *(E–H)* Results for the short timescale. Pre-retreat sessions are displayed in *(A, C, E, G)* and post-retreat are displayed in *(B, D, F, H)*, for placebo (*left*) and verum (*right*). Rare states had the highest CE_total_ regardless of timescale, condition or session. At long timescales, high-IC states had positive Δrel, whereas Δ_rel_ became negative with decreasing IC. At short timescales, this pattern was reversed, with high-IC states showing negative Δ_rel_ and low-IC states showing positive Δ_rel_. These patterns were consistent across conditions and sessions.

Thus far we have been mostly performing group level comparisons and have found timescale and network specific effects. Additionally, our global rare state analysis did not show changes related to condition or session. However, we reason that at this level stabilization constraints might be dominating what effects we can see. Therefore, we next tested whether subacute effects of condition or retreat were present in the relationship between entry and exit costs for rare states while explicitly accounting for individual differences. We fit a mixed linear model CE_out_ as a function of CE_in_, condition, and session, with random intercepts per participant. While the previous rare state analysis (Fig. 4) described topography across regions by averaging over subjects, this analysis models the relationship between CE_out_ and CE_in_ while explicitly considering individual differences. The averaged and per-subject relationships are not interchangeable. The condition effect was significant and negative for both regimes, showing that the verum group has a significantly lower pre-retreat slope than the placebo group (Table 1). In the long timescale, a session*×* condition effect on the slope emerged (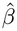= +0.014, *p* = 0.025) with the verum group’s slope showing a larger increase than placebo’s. Conceptually, for this relationship a positive slope below 1 describes sink like asymmetry where exit costs rise less quickly than entry costs. An increase in this kind of slope could therefore reflect lower sink like asymmetry and more coupling between entry and exit energy. However, this shift is small relative to the heterogeneity in individual slope changes, which span roughly -0.19 to +0.21 across the 39 subjects. Therefore, we report it as an interesting observation consistent with network findings rather than as evidence of broad energy landscape reorganization. The short timescale analysis shows no significant interaction (Table 1).

**Table 1.**
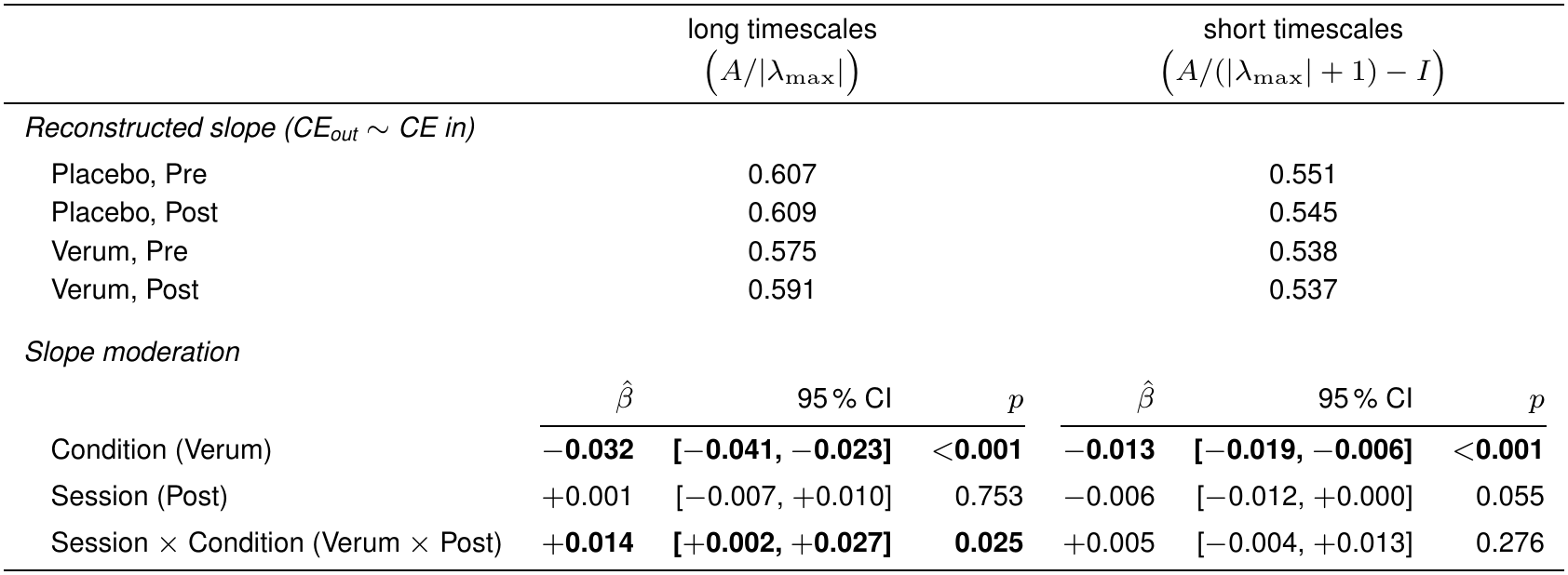
Mixed model for the analysis of CE entry and exit costs at rare functional states. Slopes (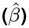) of the CE_out_ *∼*CE_in_ relationship at top-decile (*>*90th percentile) IC states, by condition and session, in both timescale regimes. Slopes are fixed-effect estimates from a linear mixed-effects model with random intercepts per subject. Reconstructed slopes are linear combinations of model coefficients relative to the reference category (placebo, pre-retreat). Slope moderation coefficients express changes in the slope relative to this reference. The long timescale regime showed a significant session*×* condition interaction in slope (*p* = 0.025), whereas the corresponding session effect on slope was not significant (*p* = 0.753). Neither the session effect on slope nor the session*×* condition interaction in slope was significant in the short timescale regime. Full model output can be found in Supplementary Table S1. Values in bold have *p <* 0.05

|  | long timescales<br>(*A*/\|λ<sub>max</sub>\|) |  |  | short timescales<br>(*A*/(\|λ<sub>max</sub>\| + 1) − *I*) |  |  |
|---|---|---|---|---|---|---|
| *Reconstructed slope (CE<sub>out</sub> ∼ CE in)* |  |  |  |  |  |  |
| Placebo, Pre | 0.607 |  |  | 0.551 |  |  |
| Placebo, Post | 0.609 |  |  | 0.545 |  |  |
| Verum, Pre | 0.575 |  |  | 0.538 |  |  |
| Verum, Post | 0.591 |  |  | 0.537 |  |  |
| *Slope moderation* |  |  |  |  |  |  |
|  | β̂ | 95 % CI | *p* | β̂ | 95 % CI | *p* |
| Condition (Verum) | −**0.032** | **[−0.041, −0.023]** | <**0.001** | −**0.013** | **[−0.019, −0.006]** | <**0.001** |
| Session (Post) | +0.001 | [−0.007, +0.010] | 0.753 | −0.006 | [−0.012, +0.000] | 0.055 |
| Session × Condition (Verum × Post) | +**0.014** | **[+0.002, +0.027]** | **0.025** | +0.005 | [−0.004, +0.013] | 0.276 |

### Psychometric correlations

Because individual responses to psychedelics are so varied we also asked whether the observed control energy effects could be related to measures of subjective experience and psychological characteristics. To do this, we computed Spearman correlations within each condition between ΔCE, calculated at both the whole-brain and network levels, and nine psychometric scales drawn from the parent study (EBI, PIS, MEQ, NADA-S, TMS-13, SOCS-S, SOCS-O, FMI Δ, NADA-T). No correlation survived FDR-BH correction at either the whole-brain or network level in either timescale regime. Thus, in this sample we did not identify robust associations between the observed control energy effects and these psychometric measures. Per-cell results, including raw *p*-values, are reported in SI Fig. S6–S7.

## Discussion

This exploratory analysis sought to investigate subacute effects of a DMT+harmine meditation retreat on the brain’s control energy landscape. Subacute effects are commonly reported in literature but are not as widely or formally studied as acute effects (28, 29). Additionally, following the finding that, acutely, psychedelics lower transition energy to traverse functional states (22) we consider it important to study this window of potential effects to understand psychedelics and their therapeutic potential. Most studies either focus on psilocybin, or group psychedelics into a broad category (43). The DMT+harmine formulation used here intends to emulate the effects of the psychedelic plant preparation Ayahuasca (37). Previous connectivity work studying subacute effects of Ayahuasca found increased connectivity within SN, decreased connectivity within DMN and increased connectivity between SN and DMN (32). However, the authors of this current dataset (33), found mixed connectivity effects, mainly on VN, SN, and attention networks, and no sustained cortical gradient disruption after the retreat (6).

Control energy quantifies the minimum input energy required to drive a transition between states given a specific structure. Changes in CE therefore reflect changes in the modeled energetic requirements of transitions, but do not necessarily directly indicate changes in neural activity and functional connectivity. We studied changes in control energy globally, across networks, and within rare states. The general pattern shows no systematic shift in control energy distribution at the transition level consistent with the absence of sustained cortical gradient disruption previously reported in this dataset (6). Average control energy was significantly higher in both groups after the retreat in the long timescale regime only and group specific effects remained at the network and state level. Importantly, results are different between the two timescale regimes, which act as different lenses on the propagation of activity across the connectome. Here, we refer to these as the short and long stabilization regimes to distinguish them from the fast and slow intrinsic dynamics of cortical networks. Although the stabilization regimes do not directly represent intrinsic cortical timescales, the networks showing effects under each regime aligned with their expected positions along the established hierarchy of intrinsic dynamics (38–40). This suggests that considering different stabilization regimes may reveal network-specific effects that could be obscured by a single stabilization choice, and motivates taking this into consideration when applying network control theory in future studies.

### Overall distribution of control costs for typical transitions is largely stable, with timescale-specific changes in mean control energy

Studies into the control energy landscape during the acute phase of psychedelics have reported a flattened energy landscape reflected in lower transition energies. We reasoned that if some aspect of that flattened landscape persisted into the subacute period it might manifest as a redistribution of transition energy costs rather than only at their mean. Therefore, we first examined distributions of CE at the transition level between session 1 and session 2 for all subjects and found no systematic distributional shift, where any significant effects were highly individual and followed no specific pattern. This argues against a broad subacute redistribution of the CE landscape and aligns with finding no cortical gradient disruptions (6). Notably, averaging across transitions showed a significant increase in mean CE in both groups under the long stabilization (Fig. 2C), while the short one showed a small, nonsignificant decrease (Fig. 2D). The shared effect under the long timescale regime is more consistent with a change associated with the retreat or other factors shared between conditions, rather than a psychedelic effect. In fact, there is evidence on the significant overlap of effects of meditation and psychedelics which has led to interest in their potential synergy (7), so the potential presence of retreat effects is not unexpected. However, functional connectivity (FC) results in this dataset found distinct patterns with decreased FC between networks due to meditation and increased FC within- and between-networks due to DMT+harmine (6). Ultimately CE and FC are different measures and can capture different properties so changes in one do not necessarily imply changes in the other. Future work could explore how changes in these metrics relate to each other. Finally, despite group patterns, individual CE trajectories are highly heterogeneous. This is consistent with inter-individual variability and subjectivity in both acute (3) and subacute (28) studies of psychedelic effects and is a well known challenge of the field.

### Network effects are stabilization specific and align with timescale hierarchies of intrinsic cortical dynamics

Although we found no systematic change in CE distributions, the concentration of mean CE changes in the long stabilization regime and network specific results in other psychedelic studies (14) led us to wonder whether subacute effects were specific to certain networks. Interestingly, the significant effects depended on the stabilization and they aligned with the hierarchy of intrinsic cortical dynamics, where, higher-order networks have been shown to exhibit slower timescales of intrinsic cortical activity than unimodal networks (40). Under the long stabilization, the verum group showed increased control energy in the CEN, with a similar but non-significant pattern in the DMN in both groups. On the other hand, in the short stabilization we saw decreased control energy in the VN. These effects are consistent across fine, middle, and coarse versions of the Schaefer parcellation (SI Figs. S4–S5). Although the stabilization regimes do not directly represent intrinsic cortical timescales, this pattern suggests that the choice of stabilization may highlight effects in networks with different dynamical properties.

Notably, these networks overlap with those showing CE reductions during an analysis on acute effects of DMT. Singleton et al. reported reductions in the visual, frontoparietal, and default mode networks, with frontoparietal and default mode effects stronger earlier after injection and visual-network effects more prominent later (23). Although there are methodological differences, this overlap suggests that similar networks may be implicated in acute and subacute DMT-related changes in CE, despite differences in the direction of effects.

The interpretation of these changes in relation to functional connectivity (FC) is less straightforward, as CE quantifies the control input required to drive transitions under a structural model and does not map directly onto FC. Nevertheless, FC findings provide a useful point of comparison, specially because there are few subacute analyses. For example, one study reported decreased within-network functional connectivity in the CEN at *∼*one week post-psilocybin (30). Reduced connectivity is one plausible explanation for increased control energy, but no corresponding CEN-specific connectivity decrease was observed in the present dataset (6), so increase in CEN control energy cannot be readily explained by reduced within-connectivity. The DMN increase is less conclusive, as it did not survive FDR correction and was observed across both groups. Nonetheless, the trend appears consistently across parcellations (SI Fig. S3A-B) so it is presented as an interesting observation. As for previous analyses of this network, psychedelics have been consistently associated with reduced DMN activity. In the acute stage studies have found decreased within-network functional connectivity, and decoupling between key hubs such as the medial prefrontal cortex (mPFC) and posterior cingulate cortex (PCC) (36, 44), a subacute study of Ayahuasca effects also found decreased PCC connectivity within the DMN (32). Comparison of meditators to controls has also shown decreased activity in mPFC and PCC in meditators (45). However, corresponding within-DMN connectivity decreases were not evident in this dataset either (6).

The decrease in control energy in the VN in the verum group under the short stabilization aligns with DMT’s characteristic heavy engagement of the visual system and the CE reductions reported by Singleton et al. (23). Additionally, the present dataset found functional connectivity effects in the VN in the verum group, mainly increased within functional connectivity and between VN and SN and between VN and DAN. Further, 5-HT_2*A*_ receptors have medium-high expression in areas of the VN (46). These findings build upon evidence showing the VN’s implication in subacute effects following DMT.

Together, these results point to network-specific subacute effects rather than uniform change in CE across the brain. Additionally, these findings are consistent with our hypothesis that network-level effects could emerge according to intrinsic timescale hierarchies, where sensory networks like the VN show short timescale effects while higher-order network effects such as in the DMN and CEN appear over a long timescale. This supports the idea that the choice of stabilization regime can highlight specific aspects of network dynamics.

### Rare state specific energy landscape is affected differently across stabilization regimes

Given highly specific effects thus far, and previous hypotheses relating psychedelic effect to an ease of transition to more states and widened repertoire (9, 22, 23), we finally wondered if subacute psychedelic effects might preferentially affect rare high energy states. The mixed-effects analysis provided limited evidence that the retreat altered the energetic organization of rare states. Under the long stabilization, the session-by-condition interaction showed a small positive increase in the slope between entry and exit costs in the verum group. As explained previously, an increase in a positive but less than 1 slope, could be interpreted as less sink like asymmetry because it reflects exit costs changing more similarly to entry costs than before. This does not necessarily confirm flattening but is consistent with it. The specificity of this result to the long stabilization regime is consistent with our network-level findings, where significant effects were also observed under long stabilization in the CEN, with a similar but nonsignificant trend in the DMN. However, the slope shift was small relative to the substantial between-subject variability, with individual changes ranging from –0.19 to +0.21. No corresponding interaction was observed under short stabilization, where the marginal session effect was primarily driven by placebo and did not reach significance. Thus, this remains a descriptive finding rather than strong evidence of lasting landscape reorganization.

Separately, rare states consistently required more total control energy to enter and exit than more common states, consistent with previous work (25). The relative costs of entry and exit, however, reversed across stabilization regimes. Under short stabilization, entering rare states was more costly than leaving them, whereas under long stabilization, leaving them was more costly. This pattern was consistent across groups and sessions, suggesting that the reversal primarily reflects the dynamical assumptions imposed by stabilization rather than a retreat-related effect. The reversal illustrates how stabilization choices shape the control-energy landscape, an important consideration given that different stabilization approaches need not be interchangeable (47). We return to the implications of this methodological choice in the following section.

### Stabilization as a methodological choice associated to timescales

Methods to stabilize the structural connectivity matrix are not typically treated as a methodological choice, with most papers using a “standard” formula that ensures that all modes decay in the absence of input (47). However, this is not the only possible assumption about brain dynamics considering that cortical activity has been shown to exhibit both attractor-like, sustaining activity and dissipative, transient regimes (48–50). Additional explanation of the motivation and formal derivation of the use of these stabilizations as differing timescale lenses can be found in Supplementary Materials. In comparing two stabilization regimes we found that the choice shaped the visible effects, with network-level findings aligning with the expected hierarchy of intrinsic cortical timescales where effects in slower networks like the CEN surface in the long timescale regime and effects in faster networks like the VN surfaced in the short timescale regime. Meanwhile, the reversal in rare-state entry and exit asymmetry across regimes was consistent across groups and sessions, suggesting that it arose primarily from the modeled dynamics rather than the retreat. Together our results show that exploring both approaches can be informative. Further investigation into the nuances of each method, and development of alternative approaches to stabilization, is warranted.

### Methodological Considerations

This work should be considered in light of the study limitations. First, sample size is small (*N* = 39), which limits statistical power. There are also methodological limitations that stem from the fact that the data was not originally collected with the intention of performing a control theoretical analysis. Because structural connectivity data was not available for individual subjects, we used a group averaged HCP structural connectivity matrix, this is an established approach in network control theory and can be appropriate when the focus is on how network structure constrains specific state transitions (22, 23, 47, 51, 52). This limited our analysis to exploring only changes in functional dynamics, forcing us to ignore potential structural differences or changes. However, this choice had the benefit of also allowing us to explore the effects of stabilization in the absence of structural heterogeneity.

Finally, subacute effects are not widely described (28) or systematically measured (29), and are highly variable (28). Therefore, further studies are required to fully piece together what effects can be reasonably attributed to the psychedelic experience, the meditation itself, or the meditation augmented by psychedelics. For instance, it would have been helpful to have a group of subjects who neither meditated nor did psychedelics, and a group of subjects who only did psychedelics, to serve as control groups. While we know that this is methodologically complex, it could provide points of comparison to better pinpoint the origins of effects. This is a potential direction for a future study in this topic.

### Conclusions and Future Work

Overall, the data and analyses presented here suggest that in the subacute stage, there is no systematic redistribution of control energy that can be attributed solely to psychedelic usage; instead, changes were specific to the timescale regimes and specific networks. Significant effects in the verum group are timescale-dependent and localized in the CEN for long timescales and the VN for short timescales. This pattern could suggest that subacute changes may be expressed differently across networks with distinct intrinsic dynamics. Although the stabilization regimes do not directly measure intrinsic cortical timescales the alignment is interesting. The rare-state analysis provided additional, tentative evidence of a verum-specific change under long stabilization, but substantial individual variability limits its interpretation as evidence of lasting landscape reorganization. We found no robust associations between CE changes and the measured psychometric scales in this dataset. Further research is needed to confirm whether these control effects are specifically related to the “afterglow” window that has been reported in the use of psychedelics (28). Given the subtlety and variability of the results, it is still possible that this effect is less about long lasting changes in brain dynamics and more related to other factors like for example a psychological openness created by the experience.

## Materials and Methods

### Data Source

This project considered resting state fMRI data from a double-blind, placebo-controlled study evaluating the effects of N,N-dimethyltryptamine (DMT) + harmine on brain connectivity during a 3-day meditation retreat (37). In this study, 40 participants that were healthy adults and experienced meditators were randomly assigned to either a placebo or verum (DMT + harmine: oromucosal doses 120mg DMT and 120mg harmine) group with 20 participants per group. Each participant was scanned one day before (session 1) and one day after (session 2) the retreat. The fMRI preprocessed data (fmriprep outputs) is openly available for download from OpenNeuro (33). Following the original functional connectivity study, subject 12 was excluded from analysis due to excessive movement (6) leaving *N* = 39 (19 Verum, 20 Placebo). No additional volume censoring was applied.

### fMRI post-processing

The preprocessed BOLD fMRI data and associated confound timeseries were obtained for each participant and session (53). All data were registered to MNI space. For each participant, parcel-level BOLD timeseries were extracted using Schaefer 100, 200 and 400 node parcellations (34). Timeseries were then cleaned with nilearn.signal.clean (54) including linear detrending, Butterworth band-pass filtering (0.01–0.1 Hz), regression of 36 nuisance signals (55), and *z*-scoring. The nuisance model comprised the 6 motion parameters together with white matter, CSF and global signal, plus their temporal derivatives, quadratic terms, and squared derivatives.

### Minimum Control Energy Computation

Network control theory (NCT) is a framework that applies concepts from graph theory and control systems engineering to understand how structure constrains and drives dynamics in a complex network. In neuroscience, it is used to model the relationship between functional brain activity and the structural connectome. In this context, we can quantify the minimal control energy required to drive the brain from one state to another. For each participant and session, we computed minimum control energy using the common linear time-invariant model (18):

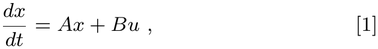

where *x* is the brain state, *A* is the structural connectivity matrix and *B* is the control input matrix. Because individual structural connectivity was not available, we used a group-average structural connectivity matrix from the Human Connectome Project (34, 56) to create an identical *A* matrix for all participants. Therefore, CE differences reflect the energetic cost of functional transitions on a canonical structural scaffold, and do not capture individual variation in structural wiring, this has been done either when individual, DTI, structural imaging is not available or specifically for group analyses (22, 23, 47, 51, 52). The control input matrix *(B)* was defined as the identity matrix, allowing for a model where control can be exerted at all brain regions. For each time step in the BOLD timeseries, we calculated the minimum control energy required for the transition from *x*_*t*_ to *x*_*t*+1_ with a time horizon of *T* = 1 using nctpy (19, 57, 58). We use node-wise contributions (*T−* 1*× N*), where each element represents the energy contribution for a given node, and the scalar total energy (*T −*1*×* 1) which is the sum across all nodes. This approach allows us to explore the control energy landscape across the whole brain and in specific networks. Main results are presented in the 200-region parcellation but were repeated in coarser (100-region) and finer (400-region) parcellations for robustness.

### Structural connectivity matrix stabilization

Stabilization of the structural connectivity matrix (*A*) is a key step in this modeling framework that offers a useful opportunity for probing brain dynamics at different timescales. The common approach in the field uses Eq. 2 in continuous-time systems (47). Following the nctpy default, we use *c* = 1.

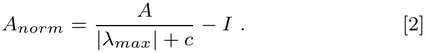

This approach ensures that all eigenvalues are negative, hence all modes decay and, in the absence of input, activity returns to zero over time (47). This choice is commonly interpreted as a way to consider dissipation due to leak currents, though it represents only one potential mathematical convention.

Here, we take the opportunity to also explore a complementary assumption, that activity does not necessarily exclusively decay between control inputs. For this, we use Eq. 3, where the spectral radius is rescaled to 1 and the dominant mode retains a positive real eigenvalue rather than the exclusively negative spectrum imposed by Eq. 2.

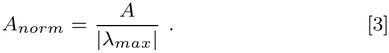

Note that this transformation, preserves eigenvectors but changes eigenvalues. This is relevant because the sign and magnitude of the eigenvalues determine whether a mode decays or expands and the rate at which it does so. Consequently, the two stabilization schemes emphasize different dynamical behaviors, with Eq. 3 emphasizing longer-timescale dynamics and Eq. 2 emphasizing shorter-timescale dynamics. Importantly, the effective integration horizon in our model remains one model time step (*T* = 1) under both schemes. Thus, we evaluate the dynamics over a finite horizon rather than considering their long-term behavior as *t→ ∞*. We report results under both regimes as complementary lenses for probing network control. A more complete explanation of how each stabilization shapes the eigenvalue distributions and, in turn, the minimum control energy calculation, is provided in the Supplementary Materials.

Because the two stabilizations impose different controllability Gramians, absolute control energy values are not directly comparable across them. Therefore, comparisons between stabilizations in this paper focus on signs and patterns rather than magnitudes.

### Total Control Energy Analysis

We explored whether there were changes across sessions and/or condition groups for the overall control energies calculated for all brain regions across the entire timeseries. In this section of the analysis we do not characterize states based on information content but analyze the obtained energies on different levels.

First, we plotted control energy distributions for each subject and each session in both groups. We performed the Kolmogorov-Smirnov tests between sessions of the same subject to observe potential distribution differences, and corrected p-values for multiple comparisons using FDR-BH. We then compared mean control energy between groups (placebo vs. verum) at each session and ΔCE computed as session 2 - session 1 using Mann-Whitney U and FDR-BH correction.

We also analyzed average control energy for each of the 7 Yeo canonical networks (35), extracted from the labels on each Schaefer parcellation. We obtained the node-wise average control energy for each subject and session, then grouped nodes according to their respective networks and averaged within-network. To determine statistical significance of differences, we used the Wilcoxon signed-rank test for session comparison, where paired values come from the same subject at two different points in time. For condition comparison, we calculated the change from session 1 to session 2 for each subject and network. We then compared the placebo and verum groups using the Mann-Whitney U test. We performed FDR-BH correction in both cases.

### Information Content Computation

To characterize the relative rarity of activation states, we computed information content for each Schaefer parcel following a probabilistic framework (25). For each participant, session, and brain region, the empirical distribution of the *z*-scored BOLD signal was estimated using Parzen-Rosenblatt kernel density estimation with a Gaussian kernel and Silverman bandwidth selection following methods from Weninger et al (25). For each parcel *i* and time point *t*, information content is defined as the negative log-probability of the observed activation magnitude:

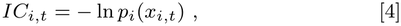

where *p*_*i*_(*x*_*i,t*_) is the estimated probability density of observing activation *x*_*i,t*_ in parcel *i*. Higher values reflect rarer activation magnitudes and lower expected probability of occurrence. The total sum across parcels defines the information content of the state (25).

Information content was computed separately for each subject and session. We used this metric as a measure of state rarity, allowing classification of brain states into low, medium, and high information states based on percentile thresholds: below the 10th percentile (bottom decile; most common states), between the 45th and 55th percentiles (middle 10%), and above the 90th percentile (top decile; rarest states). Thresholds are defined for each subject in session 1, recognizing that “high” or “low” information content can differ across subjects. This threshold definition also avoids a situation where post-retreat drug effects could shift what is considered as ‘rare’.

This separation was motivated by the hypothesis that subacute psychedelic related effects may specifically manifest in rare brain states, consistent with theories proposing a flattening of the brain energy landscape under psychedelics (22). More common states are considered in comparison to rarer states to explore associated differences.

### Information Content Stratified Analysis

We separated and labeled the bottom, middle and top IC states defining thresholds per participant at session 1. Since information content is calculated per state (*x*_*t*_), but control energy is calculated by transition (*x*_*t*_ to *x*_*t*+1_), we associate each state to a control energy required to enter it (*x*_*t−*_1 to *x*_*t*_) and another to exit it (*x*_*t*_ to *x*_*t*+1_) CE_in_ and CE_out_. The relationship between these two energies can give an idea of the general topography of the energy landscape. We plot CE_total_ = CE_in_ + CE_out_, vs the relative entry/exit difference, Δ_rel_ = (CE_out_ *−*CE_in_)*/*(CE_out_+CE_in_). For each condition, session and IC tier, CE_in_ and CE_out_ were averaged across participants and across time points within the tier to give one value per parcel (*N* = 200). Then CE_total_ and Δ_rel_ were computed from these averages. We consider CE_total_ against Δ_rel_ for session 1, session 2 and each condition group separately, with the across-parcel mean of each tier overlaid. The three tiers are discrete samples along a continuous IC scale. Each observation is one node’s CE at a single top-decile IC time point.

To formally test whether the rare-state CE landscape changed with session and condition, we fit a linear mixed-effects model with CE_out_ as the dependent variable and CE_in_, condition, session, and interactions as fixed effects, with random intercepts per participant:

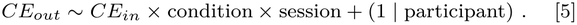

Models were fit by restricted maximum likelihood (REML) and independently estimated in Python (mixedlm from statsmodels (59)) and R (lme4 (60) with *p*-values approximated via lmerTest (61)). Both implementations produced consistent fixed-effect estimates. The *R*-fit was non-singular across all available optimizers (allFit).

### Psychometric Scales

The original study collected psychometric scales before, during, and after the retreat (37). As an exploratory follow-up to the present neural analysis, we computed Spearman correlations within each condition between Δ CE (both whole brain and per network) and nine psychometric measures: EBI, PIS, MEQ, NADA-S, TMS-13, SOCS-S, SOCS-O, FMI Δ, and NADA-T. Day 2 acute scales were collected at T+240 min (EBI, PIS, TMS-13, SOCS-S, SOCS-O) and T+360 min (MEQ, NADA-S). FMI and NADA-T were measured at baseline and post-retreat (FMI Δ = post baseline, NADA-T = post only). Correlations were FDR-BH (Benjamini–Hochberg) corrected across the full search space and were computed under both stabilizations. Because these analyses are intended as exploratory brain-experience associations, motivated by reports that psychedelic outcomes correlate with experiential intensity (62), full results are reported in the Supplementary Materials.

### Sex-stratified reproduction of results

Sex-stratified reproduction of the three main result levels (global ΔCE, network based ΔCE for the three headline networks, and rare state ΔSlope) is reported in Supplementary Table S2. Headline effects appear in both female and male subgroups in the same direction with comparable effect sizes, and no within-condition male vs. female Mann Whitney U test is significant.

### Use of AI statement

Claude Code and Claude Opus 4.8 (Anthropic) was used to support three tasks. First, Claude Code was used to debug code for data processing and analysis and suggest visualization improvements; authors reviewed, tested, and edited all AI generated code. Second, Claude Opus 4.8 was used to supplement the authors’ own literature search; the papers it suggested were manually reviewed and cited only if considered relevant and useful by the authors. Finally, Claude Opus 4.8 was used to suggest improvements on clarity, grammar, and sentence structure. The authors reviewed and edited all text, and take full responsibility for the content.

## Data and code availability

The dataset containing the preprocessed fMRI data used in this study is publicly available on Open Neuro (33). The code used to generate the analysis and figures present in this paper are available in this GitHub repository https://github.com/illano3/subacute-nct-dmt-meditation.

## References

1. I Hipólito, J Mago, FE Rosas, R Carhart-Harris, Pattern breaking: a complex systems approach to psychedelic medicine. Neurosci. Conscious. 2023, iad017 (2023).

2. GM Goodwin, et al., Single-Dose Psilocybin for a Treatment-Resistant Episode of Major Depression. New Engl. J. Medicine 387, 1637–1648 (2022) Publisher: Massachusetts Medical Society eprint: https://www.nejm.org/doi/pdf/10.1056/NEJMoa2206443.

3. LJ Mertens, et al., Methodological challenges in psychedelic drug trials: Efficacy and safety of psilocybin in treatment-resistant major depression (EPIsoDE) – Rationale and study design. Neurosci. Appl. 1, 100104 (2022).

4. TG Zaretsky, et al., The Psychedelic Future of Post-Traumatic Stress Disorder Treatment. Curr. Neuropharmacol. 22, 636–735 (2024) Publisher: Bentham Science Publishers.

5. J Richard, A Garcia-Romeu, Psychedelics in the Treatment of Substance Use Disorders and Addictive Behaviors: A Scoping Review. Curr. Addict. Reports 12, 15 (2025).

6. K Egger, et al., Meditation, psychedelics, and brain connectivity: A randomized controlled resting-state fMRI study of N,N-dimethyltryptamine and harmine in a meditation retreat. Imaging Neurosci. 3, IMAG.a.907 (2025).

7. JT Dines, Exploring the therapeutic convergence of meditation, psychedelics, and MDMA. J. Psychedelic Stud. 9, 283–293 (2025) Publisher: Akadémiai Kiadó Section: Journal of Psychedelic Studies.

8. FX Vollenweider, MF Vollenweider-Scherpenhuyzen, A Bäbler, H Vogel, D Hell, Psilocybin induces schizophrenia-like psychosis in humans via a serotonin-2 agonist action. Neuroreport 9, 3897–3902 (1998).

9. RE Daws, et al., Increased global integration in the brain after psilocybin therapy for depression. Nat. Medicine 28, 844–851 (2022) Publisher: Nature Publishing Group.

10. L Roseman, R Leech, A Feilding, DJ Nutt, RL Carhart-Harris, The effects of psilocybin and MDMA on between-network resting state functional connectivity in healthy volunteers. Front. Hum. Neurosci. 8, 204 (2014).

11. F Müller, PC Dolder, A Schmidt, ME Liechti, S Borgwardt, Altered network hub connectivity after acute LSD administration. NeuroImage: Clin. 18, 694–701 (2018).

12. DEW McCulloch, et al., Psychedelic resting-state neuroimaging: A review and perspective on balancing replication and novel analyses. Neurosci. & Biobehav. Rev. 138, 104689 (2022).

13. JS Siegel, et al., Psilocybin desynchronizes the human brain. Nature 632, 131–138 (2024) Publisher: Nature Publishing Group.

14. M Girn, et al., An international mega-analysis of psychedelic drug effects on brain circuit function. Nat. Medicine 32, 1543–1554 (2026).

15. M Girn, et al., A complex systems perspective on psychedelic brain action. Trends Cogn. Sci. 27, 433–445 (2023).

16. RL Carhart-Harris, KJ Friston, REBUS and the Anarchic Brain: Toward a Unified Model of the Brain Action of Psychedelics. Pharmacol. Rev. 71, 316–344 (2019).

17. RJ Zeifman, et al., From relaxed beliefs under psychedelics (REBUS) to revised beliefs after psychedelics (REBAS). Sci. Reports 15, 3651 (2025) Publisher: Nature Publishing Group.

18. S Gu, et al., Controllability of structural brain networks. Nat. Commun. 6, 8414 (2015).

19. TM Karrer, et al., A practical guide to methodological considerations in the controllability of structural brain networks. J. neural engineering 17, 026031 (2020).

20. Z Wu, L Huang, M Wang, X He, Development of the brain network control theory and its implications. Psychoradiology 4, kkae028 (2024).

21. WH Lee, A Rodrigue, DC Glahn, DS Bassett, S Frangou, Heritability and Cognitive Relevance of Structural Brain Controllability. Cereb. Cortex 30, 3044–3054 (2020).

22. SP Singleton, et al., Receptor-informed network control theory links LSD and psilocybin to a flattening of the brain’s control energy landscape. Nat. Commun. 13, 5812 (2022) Publisher: Nature Publishing Group.

23. SP Singleton, et al., Network control energy reductions under DMT relate to serotonin receptors, signal diversity, and subjective experience. Commun. Biol. 8, 631 (2025).

24. EJ Cornblath, et al., Temporal sequences of brain activity at rest are constrained by white matter structure and modulated by cognitive demands. Commun. Biol. 3, 261 (2020).

25. L Weninger, et al., Information content of brain states is explained by structural constraints on state energetics. Phys. Rev. E 106, 014401 (2022).

26. E Tagliazucchi, R Carhart-Harris, R Leech, D Nutt, DR Chialvo, Enhanced repertoire of brain dynamical states during the psychedelic experience. Hum. Brain Mapp. 35, 5442–5456 (2014).

27. Z Yu, et al., Alterations in brain network connectivity and subjective experience induced by psychedelics: a scoping review. Front. Psychiatry 15, 1386321 (2024).

28. R Evens, ME Schmidt, T Majić, TT Schmidt, The psychedelic afterglow phenomenon: a systematic review of subacute effects of classic serotonergic psychedelics. Ther. Adv. Psychopharmacol. 13, 20451253231172254 (2023).

29. T Majić, et al., The Afterglow Inventory (AGI): Validation of a new instrument for measuring subacute effects of classic serotonergic psychedelics. J. Psychopharmacol. 39, 474–488 (2025).

30. DEW McCulloch, et al., Lasting effects of a single psilocybin dose on resting-state functional connectivity in healthy individuals. J. Psychopharmacol. (Oxford, England) 36, 74–84 (2022).

31. RL Carhart-Harris, et al., Psilocybin for treatment-resistant depression: fMRI-measured brain mechanisms. Sci. Reports 7, 13187 (2017).

32. L Pasquini, F Palhano-Fontes, DB Araujo, Subacute effects of the psychedelic ayahuasca on the salience and default mode networks. J. Psychopharmacol. 34, 623–635 (2020) Publisher: SAGE Publications Ltd STM.

33. K Egger, M Scheidegger, DMT-HAR-MED: Effects of DMT and harmine during meditation: Functional MRI dataset (2025) [Dataset].

34. A Schaefer, et al., Local-Global Parcellation of the Human Cerebral Cortex from Intrinsic Functional Connectivity MRI. Cereb. Cortex 28, 3095–3114 (2018).

35. BT Thomas Yeo, et al., The organization of the human cerebral cortex estimated by intrinsic functional connectivity. J. Neurophysiol. 106, 1125–1165 (2011).

36. L Smigielski, M Scheidegger, M Kometer, FX Vollenweider, Psilocybin-assisted mindfulness training modulates self-consciousness and brain default mode network connectivity with lasting effects. NeuroImage 196, 207–215 (2019).

37. D Meling, et al., Meditating on psychedelics. A randomized placebo-controlled study of DMT and harmine in a mindfulness retreat. J. Psychopharmacol. 38, 897–910 (2024) Publisher: SAGE Publications Ltd STM.

38. U Hasson, E Yang, I Vallines, DJ Heeger, N Rubin, A Hierarchy of Temporal Receptive Windows in Human Cortex. J. Neurosci. 28, 2539–2550 (2008).

39. JD Murray, et al., A hierarchy of intrinsic timescales across primate cortex. Nat. Neurosci. 17, 1661–1663 (2014).

40. A Wolff, et al., Intrinsic neural timescales: temporal integration and segregation. Trends Cogn. Sci. 26, 159–173 (2022).

41. V Menon, Large-scale brain networks and psychopathology: a unifying triple network model. Trends Cogn. Sci. 15, 483–506 (2011).

42. AV Lebedev, et al., Finding the self by losing the self: Neural correlates of ego-dissolution under psilocybin. Hum. Brain Mapp. 36, 3137–3153 (2015).

43. JS Aday, CM Mitzkovitz, EK Bloesch, CC Davoli, AK Davis, Long-term effects of psychedelic drugs: A systematic review. Neurosci. & Biobehav. Rev. 113, 179–189 (2020).

44. F Palhano-Fontes, et al., The Psychedelic State Induced by Ayahuasca Modulates the Activity and Connectivity of the Default Mode Network. PLOS ONE 10, e0118143 (2015).

45. JA Brewer, et al., Meditation experience is associated with differences in default mode network activity and connectivity. Proc. Natl. Acad. Sci. 108, 20254–20259 (2011).

46. V Beliveau, et al., A High-Resolution In Vivo Atlas of the Human Brain’s Serotonin System. The J. Neurosci. 37, 120–128 (2017).

47. L Parkes, et al., A network control theory pipeline for studying the dynamics of the structural connectome. Nat. protocols 19, 3721–3749 (2024).

48. M Khona, IR Fiete, Attractor and integrator networks in the brain. Nat. Rev. Neurosci. 23, 744–766 (2022).

49. G Deco, VK Jirsa, Ongoing Cortical Activity at Rest: Criticality, Multistability, and Ghost Attractors. J. Neurosci. 32, 3366–3375 (2012).

50. BAW Brinkman, et al., Metastable dynamics of neural circuits and networks. Appl. Phys. Rev. 9, 011313 (2022).

51. AI Luppi, et al., Contributions of network structure, chemoarchitecture and diagnostic categories to transitions between cognitive topographies. Nat. Biomed. Eng. 8, 1142–1161 (2024).

52. JZ Kim, et al., Inferring intrinsic neural timescales using optimal control theory. Nat. Commun. 16, 11639 (2025).

53. O Esteban, et al., fMRIPrep: a robust preprocessing pipeline for functional MRI. Nat. Methods 16, 111–116 (2019).

54. A Abraham, et al., Machine learning for neuroimaging with scikit-learn. Front. Neuroinformatics 8 (2014).

55. TD Satterthwaite, et al., An improved framework for confound regression and filtering for control of motion artifact in the preprocessing of resting-state functional connectivity data. NeuroImage 64, 240–256 (2013).

56. S Lariviére, et al., The ENIGMA Toolbox: multiscale neural contextualization of multisite neuroimaging datasets. Nat. Methods 18, 698–700 (2021) Publisher: Nature Publishing Group.

57. L Parkes, JZ Kim, J Stiso, nctpy: Network Control Theory for Python. (2024).

58. JZ Kim, DS Bassett, Linear Dynamics and Control of Brain Networks in Neural Engineering, ed. B He. (Springer International Publishing, Cham), pp. 497–518 (2020).

59. S Seabold, J Perktold, Statsmodels: Econometric and Statistical Modeling with Python. SciPy 2010 (2010).

60. D Bates, M Mächler, B Bolker, S Walker, Fitting Linear Mixed-Effects Models Using lme4. J. Stat. Softw. 67, 1–48 (2015).

61. A Kuznetsova, PB Brockhoff, RHB Christensen, lmerTest Package: Tests in Linear Mixed Effects Models. J. Stat. Softw. 82, 1–26 (2017).

62. B Romeo, et al., The intensity of the psychedelic experience is reliably associated with clinical improvements: A systematic review and meta-analysis. Neurosci. & Biobehav. Rev. 172, 106086 (2025).

