## Supplementary Information for "Exploring subacute effects of a DMT and harmine meditation retreat on the brain’s control energy landscape"

#### This PDF file includes:

Supporting text
Figs. S1 to S7
Tables S1 to S2
SI References

### Supporting Information Text

**Mathematical and methodological explanation of timescale lens approximation.** This section will expand upon the mathematical basis for the use of different $A$ matrix stabilization regimes to emphasize activity occurring at different timescales. Current field convention employs equation 1, usually with a default value of $c = 1$ (1). This approach makes all eigenvalues negative which means that they all decay in the absence of input and so over time, activity will go to zero (1).

$$A_{norm} = \frac{A}{|\lambda_{max}| + c} - I \tag{1}$$

In traditional control theory, analyses often consider the dynamics of a system at $t \to \infty$ because the long term stability is of great importance. This is part of what motivates the desire to make all eigenvalues negative, as in equation 1. The other part is an argument about leak currents in the brain and how activity does not grow to infinity. However, we always interrogate the system at a time horizon of $T = 1$, not infinity, and arguments for the existence of self-sustained activity in the brain can also be made (2–4). To address this, we also employ the normalization approach described by equation 2 in which the spectral radius is rescaled to 1. This leaves the dominant mode with a positive real eigenvalue, and thus activity is not necessarily forced to decay in the absence of input.

$$A_{norm} = \frac{A}{|\lambda_{max}|} \tag{2}$$

These forms of normalization differ only in the scaling of their eigenvalues, and thus the rate of decay or growth in the system. Neither operation rotates an eigenbasis, so the two share eigenvectors exactly and differ only in their eigenvalues. Each mode is an eigenvector–eigenvalue pair $(v_i, \lambda_i)$ in which $v_i$ is a spatial pattern and $\lambda_i$ the rate at which it grows or decays. Since $A$ is symmetric, all eigenvalues are real and no mode oscillates. Free evolution scales each mode by $e^{\lambda_i T}$, so each has a characteristic time $\tau_i = 1/|\lambda_i|$ over which its pattern decays ($\lambda < 0$) to $1/e$ of its amplitude or expands by a factor of $e$ ($\lambda > 0$).

When we use this model to analyze brain activity as a controllable network, we calculate control energy every one model time step $T$. It is important to highlight that this is not necessarily time as understood by $TR$ in the fMRI collection protocol or time as understood as seconds passed from a certain moment. At this point in the modeling we are in “model time", and so $T$ is simply a step whose length is determined by our time horizon, which is always 1 in our case. The concern of activity reaching infinity or zero does not occur at these short time steps. We never explore the system as $t \to \infty$; instead we evaluate control energy over a single transition, and the Gramian is integrated over a finite horizon. So the free response is never propagated for more than one step.

In what we call the short timescale regime (equation 1), characteristic times shorten and are mostly compressed to one model step $T$. This is an uneven process, where already fast modes change relatively little while slow modes are much more compressed as shown below:

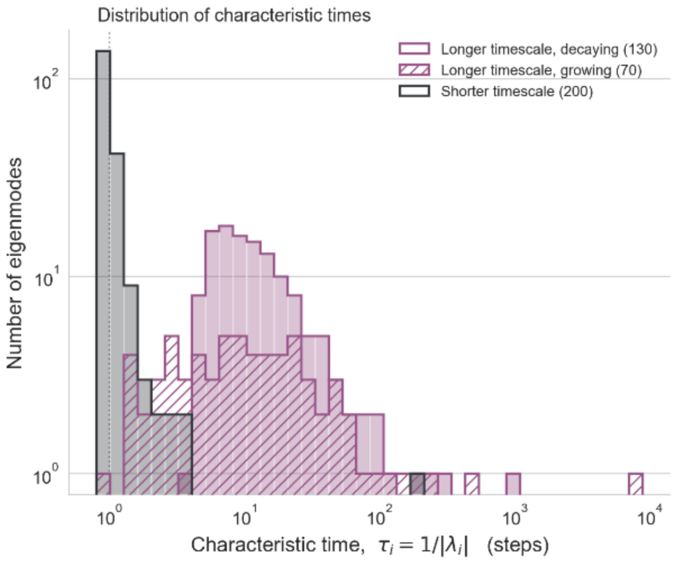


Distribution of characteristic times for modes under both stabilization regimes. The distribution for the 200 decaying modes of the short timescale regime are depicted in gray. Distribution for modes under the longer timescale regime are shown in pink. Decaying modes are shown in solid color while expansive modes are shown in stripes. The longer timescale modes occupy a much broader distributon of characteristic times while the short timescale modes are mostly compressed to one model step.

Conversely, in the long timescale regime (equation 2) we don’t assume that all modes must decay. This reduces the compression and allows for the existence of some expanding modes ((34/100 (34.0%), 70/200 (35.0%) and 142/400 (35.5%) at

Schaefer-100, 200 and 400 respectively). This is what allows us to question the system at different timescales. This lens is more evident when we compare median characteristic times ($\tau_i$) for decaying modes across parcellations, which are 10.57, 10.99 and 11.66 steps under equation 2, compared to 0.95, 0.96 and 0.96 under equation 1 for Schaefer 100, 200 and 400 respectively. Under equation 2, roughly a third of modes are not required to decay at all, and those that do decay do so roughly an order of magnitude more slowly. Therefore, the same transition is evaluated over a substantially longer effective window. These two regimes act as complimentary lenses to a same system.

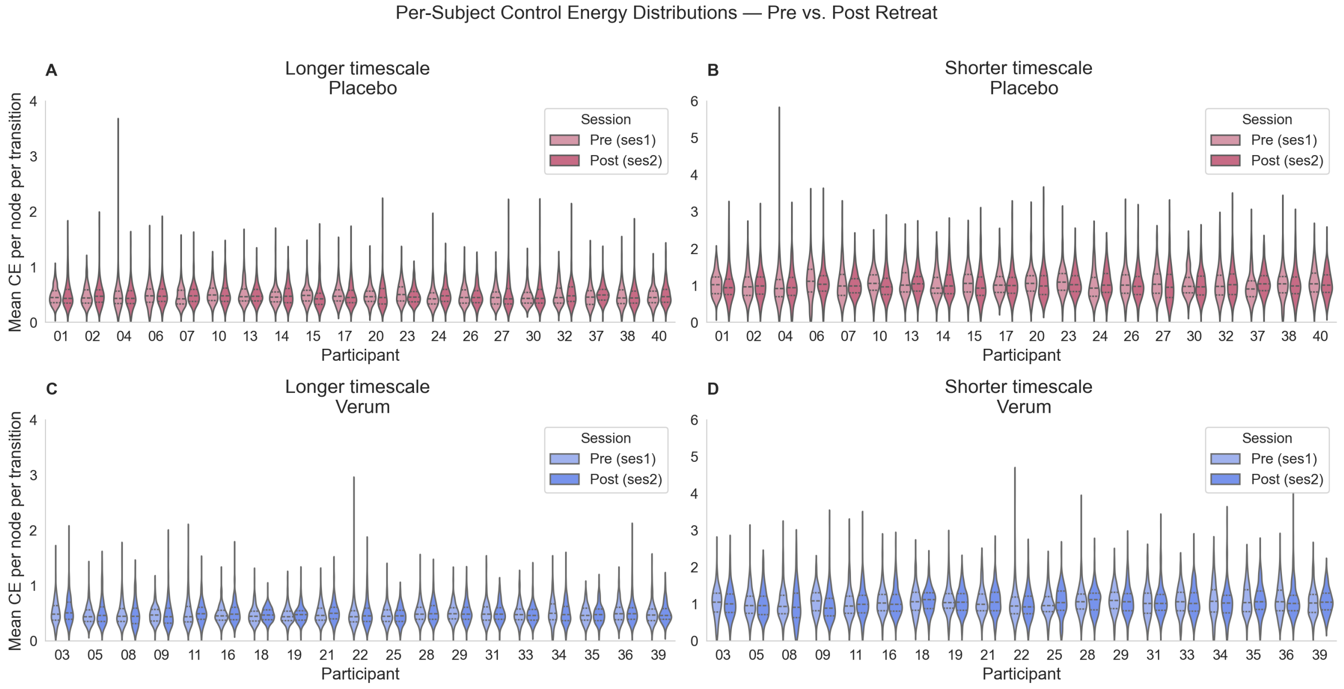


**Fig. S1.** Each violin represents the distribution of mean control energy (CE) per node across all time-point transitions for a single participant. Light and dark shading correspond to pre-retreat (Session 1) and post-retreat (Session 2) recordings, respectively. Panels A and C show results under the Longer timescale stabilization while panels B and D show the Shorter timescale stabilization. Placebo participants are shown in pink (top row) and verum participants in teal (bottom row).

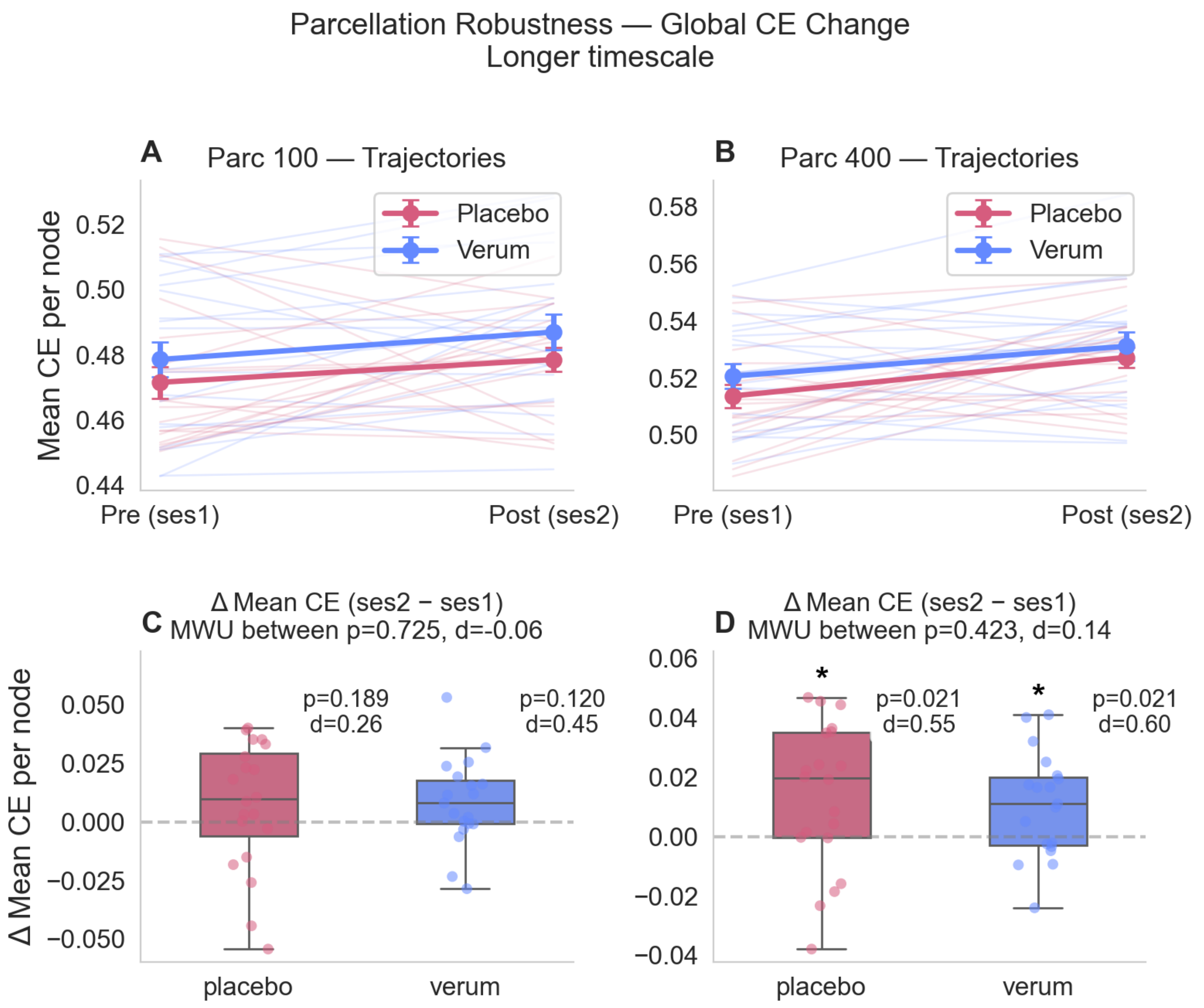


**Fig. S2.** Mean CE trajectories and mean CE change from session 1 to session 2 for 100 and 400 Schaefer parcellations in the Longer timescale stabilization.

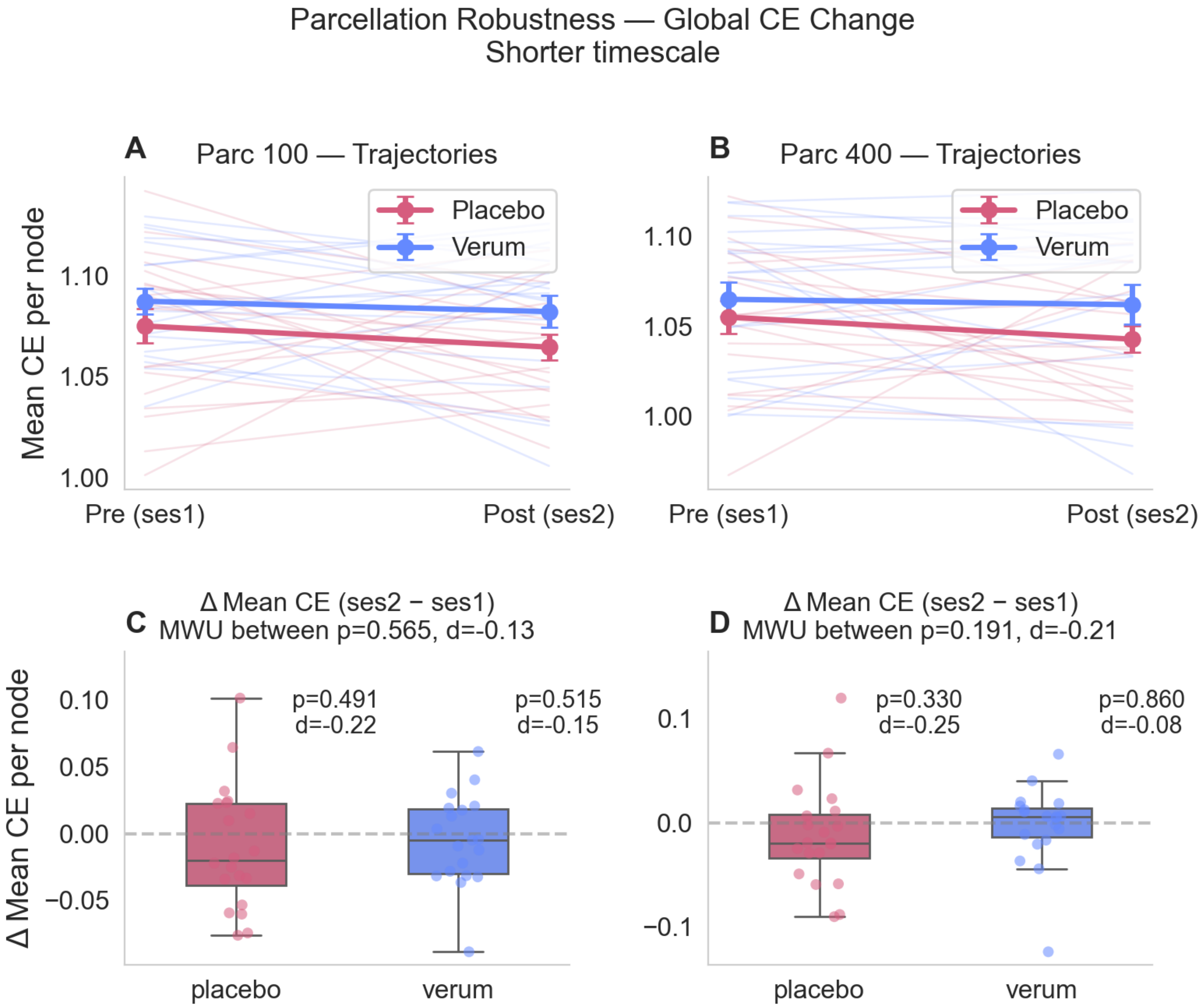


**Fig. S3.** Mean CE trajectories and mean CE change from session 1 to session 2 for 100 and 400 Schaefer parcellations in the Shorter timescale stabilization.

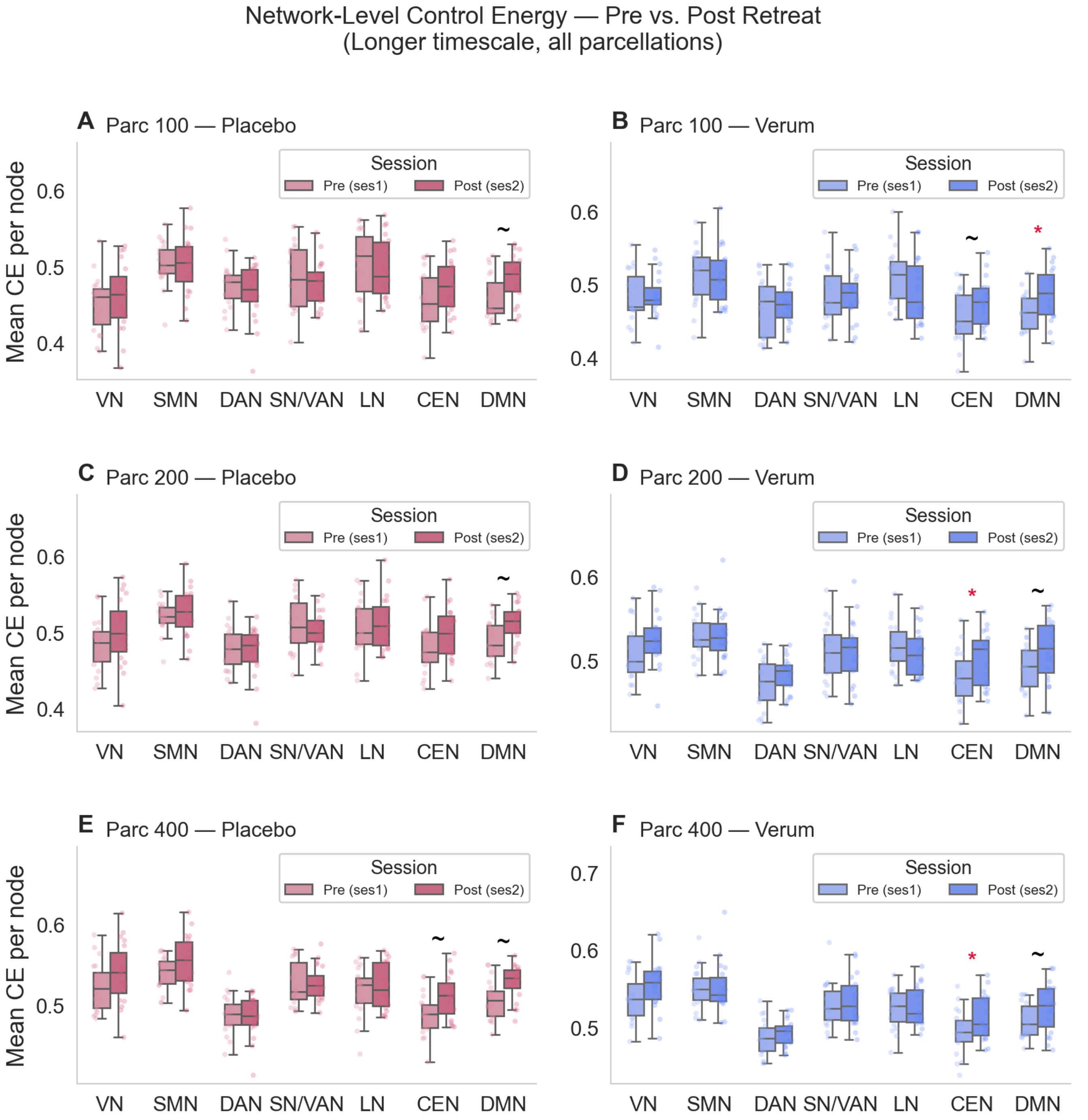


**Fig. S4.** Longer timescale stabilization mean CE for 100, 200 and 400 Schaefer parcellations divided by Yeo-7 network. Red * indicates statistically significant difference between session 1 and session 2 that survived FDR correction for multiple comparisons. indicates statistical significance that did not survive FDR correction for multiple comparisons.

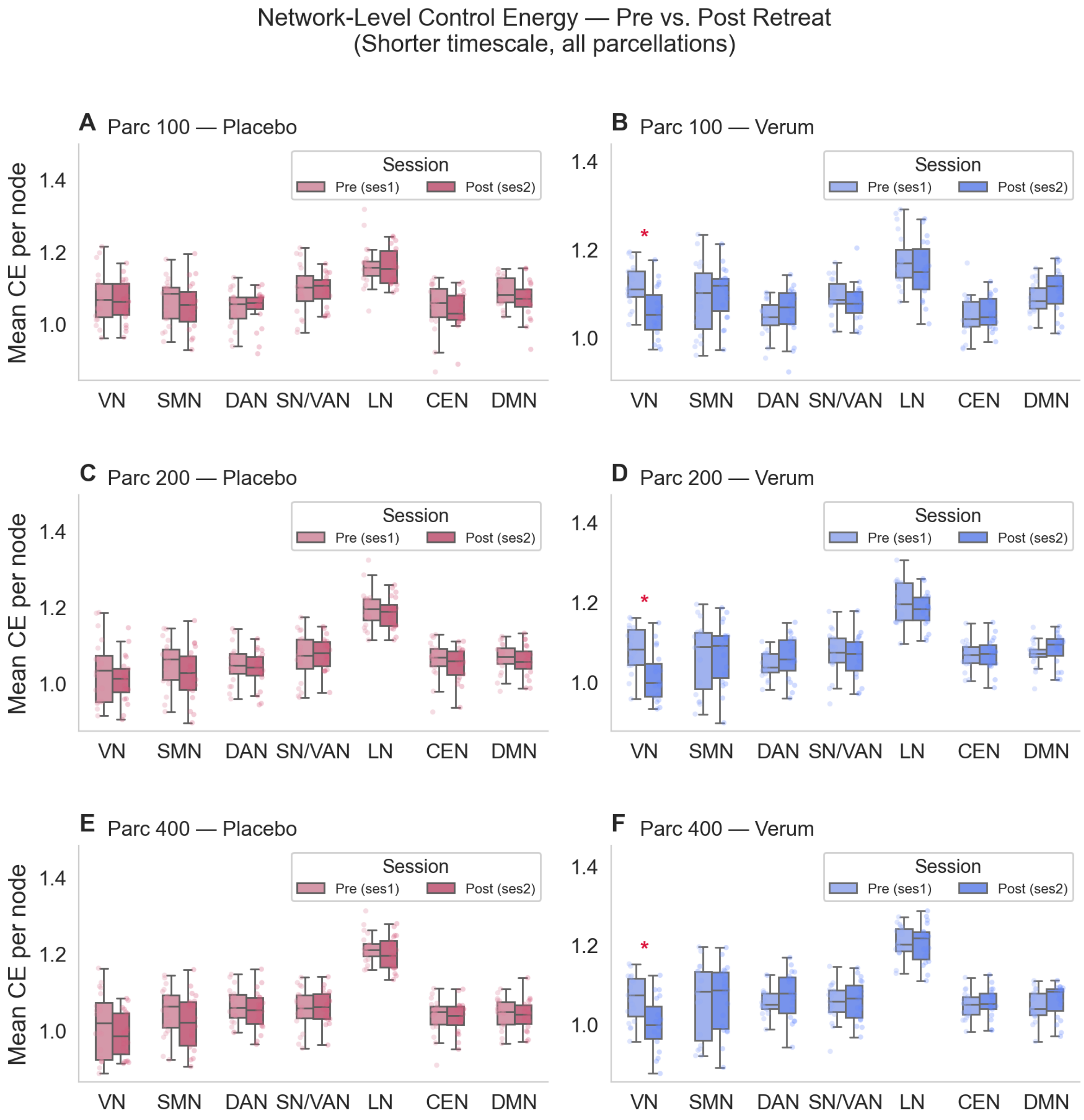


**Fig. S5.** Shorter timescale stabilization mean CE for 100, 200 and 400 Schaefer parcellations divided by Yeo-7 network. Red * indicates statistically significant difference between session 1 and session 2 that survived FDR correction for multiple comparisons.   indicates statistical significance that did not survive FDR correction for multiple comparisons.

**Table S1. Full mixed-model output for the analysis of CE entry and exit costs at rare functional states. Complete fixed-effects table for the linear mixed-effects model ($CE_{out} \sim CE_{in} \times$ condition $\times$ session, random intercepts per subject). All terms are relative to the reference category (placebo, pre-retreat). Values in bold have $p < 0.05$.**

| | long timescales $\left(A/\lvert\lambda_{\max}\rvert\right)$ | | | short timescales $\left(A/(\lvert\lambda_{\max}\rvert+1)-I\right)$ | | |
|---|---|---|---|---|---|---|
| | $\hat{\beta}$ | 95 % CI | $p$ | $\hat{\beta}$ | 95 % CI | $p$ |
| Intercept | **+0.370** | **[+0.350, +0.389]** | **<0.001** | **+0.467** | **[+0.439, +0.495]** | **<0.001** |
| $CE_{in}$ (Placebo, Pre) | **+0.607** | **[+0.601, +0.614]** | **<0.001** | **+0.551** | **[+0.547, +0.555]** | **<0.001** |
| Condition (Verum) | +0.003 | [−0.025, +0.031] | 0.829 | +0.007 | [−0.033, +0.047] | 0.738 |
| Session (Post) | **+0.035** | **[+0.025, +0.045]** | **<0.001** | −0.012 | [−0.030, +0.005] | 0.177 |
| $CE_{in} \times$ Condition (Verum) | **−0.032** | **[−0.041, −0.023]** | **<0.001** | **−0.013** | **[−0.019, −0.006]** | **<0.001** |
| $CE_{in} \times$ Session (Post) | +0.001 | [−0.007, +0.010] | 0.753 | −0.006 | [−0.012, +0.000] | 0.055 |
| Condition $\times$ Session | **−0.026** | **[−0.041, −0.012]** | **<0.001** | +0.021 | [−0.004, +0.047] | 0.102 |
| $CE_{in} \times$ Session $\times$ Condition (Verum $\times$ Post) | **+0.014** | **[+0.002, +0.027]** | **0.025** | +0.005 | [−0.004, +0.013] | 0.276 |

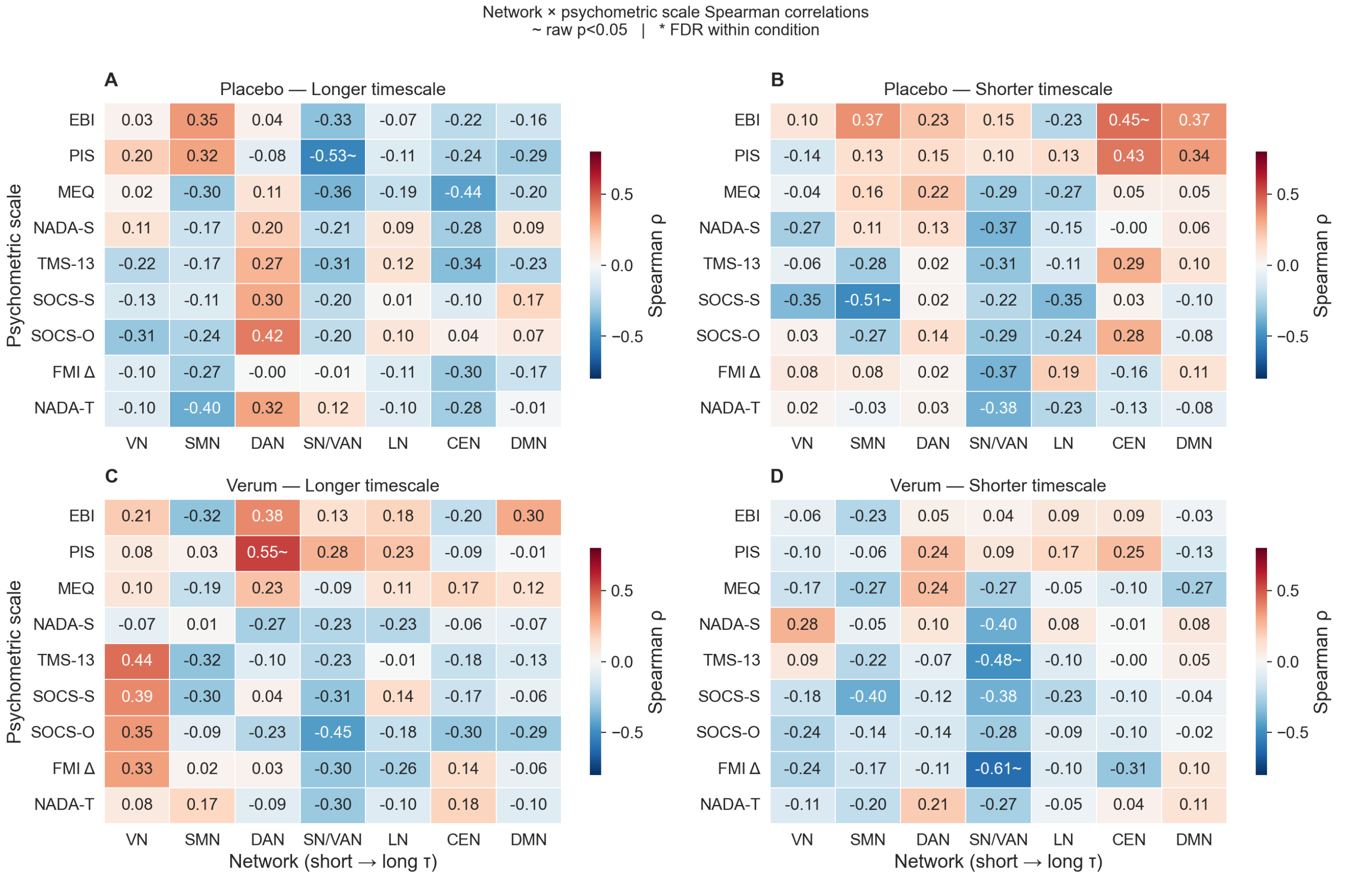


**Fig. S6.** Spearman correlation of Δ mean CE by network and score in psychometric scale. No correlation survives FDR-BH correction for multiple comparisons. There are also no consistent correlation patterns for specific network x psychometric scale combinations.

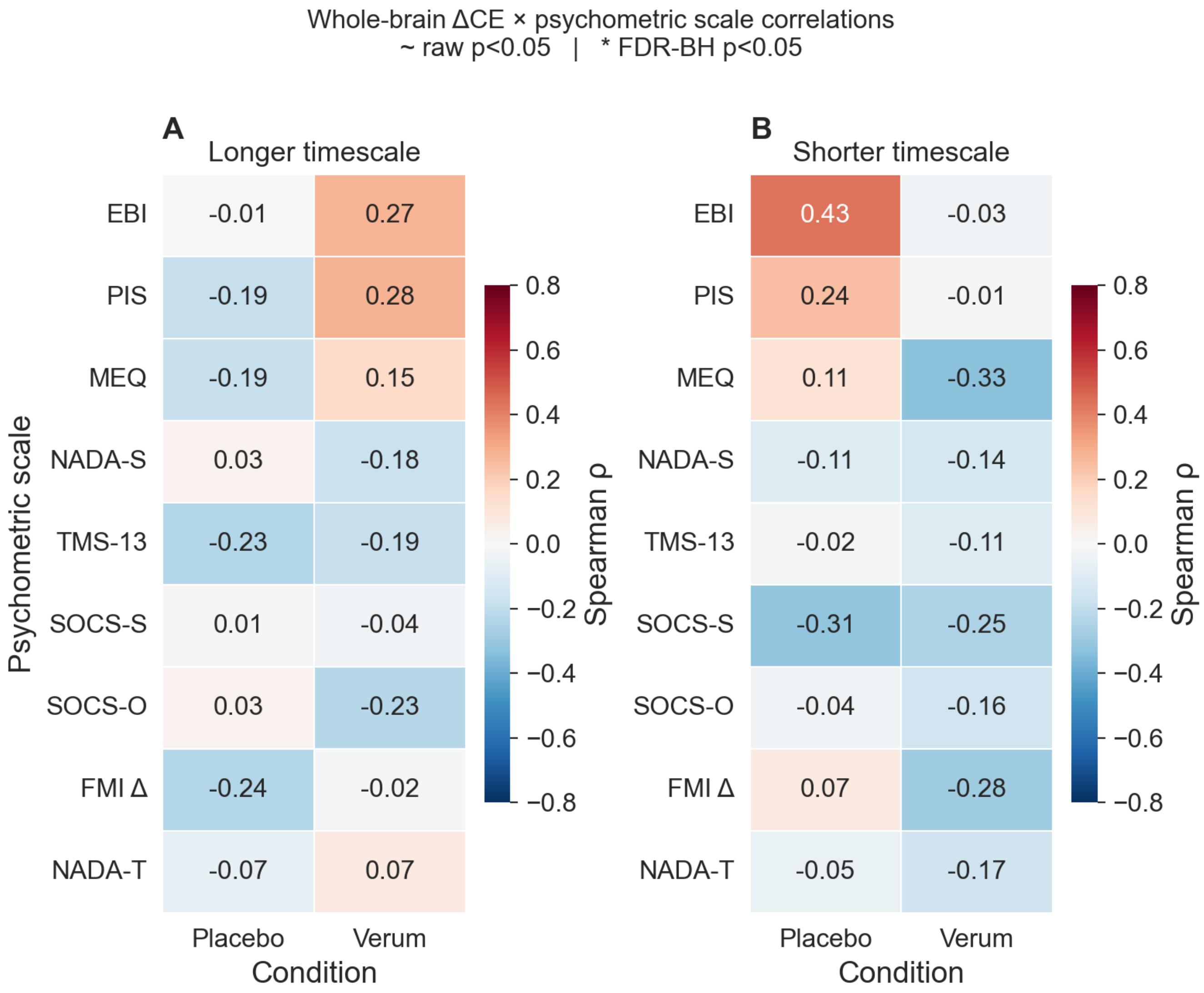


**Fig. S7.** Spearman correlations between whole brain △ mean CE and psychometric scales. No correlation is statistically significant raw or after FDR-BH multiple comparisons correction.

**Table S2. Sex-stratified reproduction of the two main results Global $\triangle$CE and network-based $\triangle$CE for the three headline networks, split by sex (F: 17 | M: 22). Headline effects appear in both female and male subgroups in the same direction with comparable effect sizes, and no within-condition male vs. female Mann–Whitney $U$ test is significant. Values in bold have $p < 0.05$.**

| Result level | Stabilization | Network | Cond. | All ($d_z$, $p$) | F-only ($d_z$, $p$) | M-only ($d_z$, $p$) | M vs. F ($p$) |
|---|---|---|---|---|---|---|---|
| Global $\triangle$CE | Longer timescale | — | Placebo | **+0.46 (0.044)** | +0.14 (0.652) | **+0.84 (0.014)** | 0.323 |
| | | | Verum | **+0.60 (0.020)** | +0.67 (0.148) | +0.52 (0.206) | 0.778 |
| | Shorter timescale | — | Placebo | −0.28 (0.154) | −0.12 (0.359) | −0.40 (0.240) | 0.649 |
| | | | Verum | −0.14 (0.709) | −0.52 (0.148) | +0.22 (0.413) | 0.091 |
| Network $\triangle$CE | Longer timescale | CEN | Placebo | +0.41 (0.143) | +0.10 (0.820) | +0.65 (0.067) | 0.224 |
| | | | Verum | **+0.83 (0.002)** | +0.67 (0.109) | **+0.93 (0.019)** | 0.840 |
| | | DMN | Placebo | **+0.60 (0.019)** | +0.37 (0.301) | +0.77 (0.083) | 0.323 |
| | | | Verum | **+0.69 (0.011)** | +0.83 (0.078) | +0.56 (0.102) | 0.545 |
| | | VN | Placebo | +0.33 (0.202) | +0.34 (0.496) | +0.31 (0.365) | 0.761 |
| | | | Verum | +0.37 (0.134) | +0.39 (0.312) | +0.34 (0.278) | 0.600 |
| | Shorter timescale | CEN | Placebo | −0.18 (0.388) | −0.08 (0.910) | −0.23 (0.465) | 0.543 |
| | | | Verum | +0.06 (0.922) | −0.20 (0.250) | +0.23 (0.465) | 0.272 |
| | | DMN | Placebo | −0.15 (0.452) | +0.07 (0.910) | −0.33 (0.320) | 0.494 |
| | | | Verum | +0.29 (0.196) | +0.06 (0.844) | +0.44 (0.175) | 0.272 |
| | | VN | Placebo | −0.23 (0.312) | −0.21 (0.570) | −0.24 (0.520) | 0.820 |
| | | | Verum | **−0.91 (0.002)** | **−1.03 (0.008)** | −0.81 (0.054) | 0.395 |

### References

1. L Parkes, et al., Using network control theory to study the dynamics of the structural connectome. *bioRxiv* p. 2023.08.23.554519 (2023).
2. M Khona, IR Fiete, Attractor and integrator networks in the brain. *Nat. Rev. Neurosci.* **23**, 744–766 (2022).
3. G Deco, VK Jirsa, Ongoing Cortical Activity at Rest: Criticality, Multistability, and Ghost Attractors. *J. Neurosci.* **32**, 3366–3375 (2012).
4. BAW Brinkman, et al., Metastable dynamics of neural circuits and networks. *Appl. Phys. Rev.* **9**, 011313 (2022).
